# *Pseudomonas aeruginosa* type IV minor pilins and PilY1 regulate virulence by modulating FimS-AlgR activity

**DOI:** 10.1101/192062

**Authors:** Victoria A. Marko, Sara L.N. Kilmury, Lesley T. MacNeil, Lori L. Burrows

## Abstract

Type IV pili are expressed by a wide range of prokaryotes, including the opportunistic pathogen *Pseudomonas aeruginosa*. These flexible fibres mediate twitching motility, biofilm maturation, surface adhesion, and virulence. The pilus is composed mainly of major pilin subunits while the low abundance minor pilins FimU-PilVWXE and the putative adhesin PilY1 prime pilus assembly and are proposed to form the pilus tip. The minor pilins and PilY1 are encoded in an operon that is positively regulated by the FimS-AlgR two-component system. Independent of pilus assembly, PilY1 is proposed to be a mechanosensory component that - in conjunction with minor pilins - triggers up-regulation of acute virulence phenotypes upon surface attachment. Here, we investigated the link between the minor pilins and virulence. *pilW, pilX*, and *pilY1* mutants had reduced virulence towards *Caenorhabditis elegans* relative to wild type or a major pilin mutant, implying a role in pathogenicity that is independent of pilus assembly. We hypothesized that loss of specific minor pilins relieves feedback inhibition on FimS-AlgR, increasing transcription of the minor pilin operon and other members of the AlgR regulon. Reporter assays confirmed that FimS-AlgR were required for the increased expression from the minor pilin operon promoter upon loss of select minor pilins. Overexpression of AlgR or its hyperactivation via point mutation reduced virulence, and the virulence defects of *pilW*, *pilX*, and *pilY1* mutants were dependent on FimS-AlgR expression and activation. We propose that PilY1 and the minor pilins inhibit their own expression, and that loss of these proteins leads to FimS-mediated activation of AlgR and reduced expression of acute-phase virulence factors. This mechanism could contribute to adaptation of *P. aeruginosa* in chronic lung infections, as mutations in the minor pilin operon result in the loss of piliation and increased expression of AlgR-dependent virulence factors – such as alginate – that are characteristic of such infections.

**Author summary:** *Pseudomonas aeruginosa* causes dangerous infections, including chronic lung infections in cystic fibrosis patients. It uses many strategies to infect its hosts, including deployment of grappling hook-like fibres called type IV pili. Among the components involved in assembly and function of the pilus are five proteins called minor pilins that - along with a larger protein called PilY1 - may help the pilus attach to surfaces. In a roundworm infection model, loss of PilY1 and specific minor pilins delayed killing, while loss of other pilus proteins did not. We traced this effect to increased activation of the FimS-AlgR regulatory system that inhibits expression of virulence factors used to initiate infections, while positively regulating chronic infection traits such as alginate production, a phenotype called mucoidy. A disruption in the appropriate timing of FimS-AlgR-dependent virulence factor expression when select minor pilins or PilY1 are missing may explain why those pilus-deficient mutants have reduced virulence compared with others whose products are not under FimS-AlgR control. Increased FimS-AlgR activity upon loss of PilY1 and specific minor pilins could help to explain the frequent co-occurrence of the non-piliated and mucoid phenotypes that are hallmarks of chronic *P. aeruginosa* lung infections.

## Introduction

*Pseudomonas aeruginosa* is a Gram-negative opportunistic pathogen, recently listed as one of the highest priority antimicrobial-resistant threats by the World Health Organization, due to its intrinsic antibiotic resistance and recalcitrance to therapy [1]. Among its virulence factors are filamentous surface appendages called type IV pili (T4P), sophisticated biological nanomachines that are broadly distributed among bacteria and archaea [2, 3]. In *P. aeruginosa*, T4P facilitate surface and host cell adhesion, colonization, biofilm maturation, virulence, and twitching, a form of surface-associated motility facilitated by cycles of extension, adhesion, and retraction of T4P fibres [3-11]. T4P are composed of hundreds to thousands of copies of small proteins called major pilins (PilA in *P. aeruginosa*) along with the low abundance minor pilins (MPs) FimU-PilVWXE [12-16]. The MPs are encoded in a polycistronic operon with the *pilY1* gene that codes for a large ~125 kDa non-pilin protein. The operon is positively regulated by the virulence factor regulator Vfr, and the two-component system (TCS) FimS (AlgZ)-AlgR. FimS is a predicted histidine sensor kinase and AlgR is a response regulator that promotes expression of genes important for biofilms and chronic cystic fibrosis (CF) lung infections [17-21]. The N-termini of immature pilins are cleaved and methylated at the cytoplasmic face of the inner membrane (IM) by the prepilin peptidase, PilD, while PilY1 may be processed by signal peptidase 1 [22-25]. Mature pilins are polymerized into a T4P fibre via an envelope-spanning assembly machinery, where individual PilA subunits are added or removed at the platform protein, PilC, via action of the ATPases PilB and PilT, respectively [2, 26].

The MPs and PilY1 are required for T4P function in several bacterial species, including *P. aeruginosa*, *Escherichia coli*, *Neisseria meningitidis*, *N. gonorrhoeae*, and *Myxococcus xanthus* [12-15, 27-30]. PilY1 and the MPs were originally proposed to oppose pilus retraction, as some surface pili remain in *pilY1* and MP mutants when retraction is blocked via deletion of *pilT* [23, 28, 29, 31, 32]. We recently showed that deletion of the minor pseudopilins of the Xcp type II secretion system in a *pilT* background lacking the T4P MPs abolished pilus assembly, suggesting that when MPs are missing, the minor pseudopilins can prime extension, but cannot counteract retraction [24]. We also demonstrated that PilY1 and the MPs are present in sheared pili, and that the loss of PilV, PilW, PilX, or PilY1 excludes the other three components from the pilus [24]. Thus, PilVWXY1 are proposed to form a core assembly-initiation subcomplex, while FimU and PilE are thought to connect this complex to PilA. Initiation of assembly with subsequent addition of multiple PilA subunits would place the MPs at the pilus tip, with PilY1 – the largest component – at the distal position, supporting the hypothesis that PilY1 is a T4P-associated adhesin [31].

PilY1 and the MPs (and their regulators FimS-AlgR) are required for T4P biogenesis, and therefore T4P-mediated function [12-15, 17, 19]. However, recent studies hinted at more enigmatic roles of PilWXY1 in virulence. Bohn et al. [33] showed that in a non-piliated *P. aeruginosa* background, subsequent loss of *pilY1* reduced virulence in a *Caenorhabditis elegans* fast killing assay and in a mouse airway infection model, and increased resistance to killing by neutrophils. Thus, PilY1 has a role in virulence that does not require functional pili. Other studies using *C. elegans* infection models suggested that MP and *pilY1* mutants had attenuated virulence relative to WT, and in one case, to a non-piliated mutant [34-37]. Recently, Siryaporn et al. [38] showed that PilWXY1 were required for surface-activated virulence towards amoebae, while other non-piliated mutants had WT virulence. The N-terminal region of PilY1 has limited sequence similarity to the eukaryotic von Willebrand factor A (VWFa) domain, which can be deformed by shear forces [39]. In-frame deletion of this domain from PilY1 allowed normally avirulent planktonic cells to kill amoebae [38]. PilY1 was therefore proposed to be a mechanosensor, where deformation of its VWFa domain upon surface interaction led – by an as-yet unknown mechanism – to increased expression of virulence factors. One important caveat of that study was that an *algR* mutant (which also lacks PilY1 and the MPs) had WT virulence towards amoebae [38].

Deformation of PilA subunits by tensile forces acting upon surface-attached pili was also proposed as a possible way to signal attachment. Detection of partly unfolded pilins by the Pil-Chp chemotaxis system could lead to increased cyclic adenosine monophosphate (cAMP) synthesis via the CyaB adenylate cyclase [40, 41]. cAMP is bound by Vfr, a key transcription factor that promotes expression of virulence factors involved in motility, attachment, and secretion [20, 40, 41]. *fimS-algR* transcription is activated by Vfr, leading to increased transcription of *fimU*-*pilVWXY1E* [40]. PilVWXY1 were proposed to repress their own expression in an AlgR-dependent manner, as the loss of *pilV*, *pilW*, *pilX*, or *pilY1* led to elevated expression of the MP operon and *fimS-algR* [23, 33, 38, 40]. The mechanism of this putative feedback inhibition is largely uncharacterized, but was speculated to involve FimS [40].

Once expression of the MP operon is activated, extracellular PilY1 may sense surface association via its VWFa domain and transduce this information through the T4P assembly machinery [38, 40]. This signal is thought to activate an IM-localized diguanylate cyclase, SadC, to increase levels of c-di-GMP, promoting expression of genes associated with a biofilm lifestyle, while repressing early-phase virulence traits such as swarming motility [40, 42]. This model was supported by studies demonstrating that loss of *pilW*, *pilX*, or *pilY1* in a high-c-di-GMP background resulted in hyper-swarming and reduced c-di-GMP levels, as measured by liquid chromatography-mass spectrometry of extracts from surface-grown cells [39, 43]. Rodesney et al. [44] showed that c-di-GMP levels increased in response to shear forces, and that functional T4P were required for this phenomenon, further supporting this hypothesis. However, unlike *pilW*, *pilX*, and *pilY1* mutants, a *sadC* mutant had WT virulence towards amoebae, suggesting the PilWXY1-SadC pathway may be important for surface sensing, but not necessarily for surface-activated virulence [38].

Although PilY1 and the MPs clearly influence virulence, the underlying mechanism remains to be established [33-36, 38, 45]. We hypothesized that a subset of these components represses FimS activity, such that loss of *pilW*, *pilX*, or *pilY1* activates FimS-AlgR, shifting the bacteria to a less pathogenic phenotype typically associated with chronic infection. We found that *pilW*, *pilX*, and *pilY1* mutants had attenuated virulence in *C. elegans* slow killing (SK) assays compared to WT or a *pilA* mutant, and this was dependent on FimS-AlgR, because double mutants had WT virulence. Hyperactivation (via phospho-mimetic point mutation) or overexpression of AlgR alone was sufficient to attenuate virulence. Together, these data are consistent with a model where loss of PilWXY1 relieves feedback inhibition on expression of the AlgR regulon, resulting in dysregulation of virulence factors that are important for *C. elegans* pathogenesis.

## Results

### PilWXY1 are important for T4P-independent virulence in PA14 and PAO1

Specific genes in the MP operon were reported to be important for virulence in amoebae, nematodes, and mouse models, but those studies were done using different strains of *P. aeruginosa* [33-36, 38, 45]. We first sought to confirm these results in the *C. elegans* SK model, using two well-studied strains. SK assays were performed using PA14 with deletions of *pilA*, *fimU*, *pilV*, *pilW*, *pilX*, *pilY1*, or *pilE* (Fig 1A). An *E. coli* OP50 plate was included as a negative control for pathogenicity; worms began to senesce on these plates around day 7-8, consistent with published data regarding temperature-dependent effects on lifespan [46]. Given that worms at later time points were at increased risk of death due to ageing in addition to *P. aeruginosa* infection, statistical significance was assessed using the Gehan-Breslow-Wilcoxon test, which places greater weight on earlier time points [47]. A *pilA* (major pilin) mutant was slightly less virulent than WT; subsequent comparisons were made relative to *pilA,* since all mutants lack pili. *fimU* and *pilE* mutants had increased virulence relative to the *pilA* mutant, with similar virulence to WT. In contrast, *pilW*, *pilX*, and *pilY1* mutants had reduced virulence relative to the *pilA* mutant, suggesting their reduced virulence was not due to loss of functional T4P. Virulence of the *pilV* mutant was similar to the *pilA* mutant. The twitching and virulence defects of *pilW*, *pilX*, and *pilY1* mutants could be partially complemented by expression of the relevant gene *in trans* (Supplementary Fig S1). The stoichiometry of PilY1 and the MPs is important for optimal T4P function [23], which may explain the lack of full complementation. To verify that these phenotypes were not strain-specific, we tested PAO1 transposon-insertion mutants of *pilA*, *fimU*, *pilV*, *pilW*, *pilX*, *pilY1*, and *pilE* in the SK assay (Fig 1B). Similar to the results in PA14, PilWXY1 were important for T4P-independent virulence. However, the *fimU* and *pilV* mutants were also less pathogenic than *pilA*; the PA14 and PAO1 MPs are divergent (61-75% amino acid similarity), so it is possible that FimU and PilV function slightly differently in PAO1 versus PA14 [48]. To focus on genes that were generally important for virulence of *P. aeruginosa*, we undertook studies of the mechanism responsible for loss of virulence in the *pilW*, *pilX*, and *pilY1* mutants.

**Fig 1.**
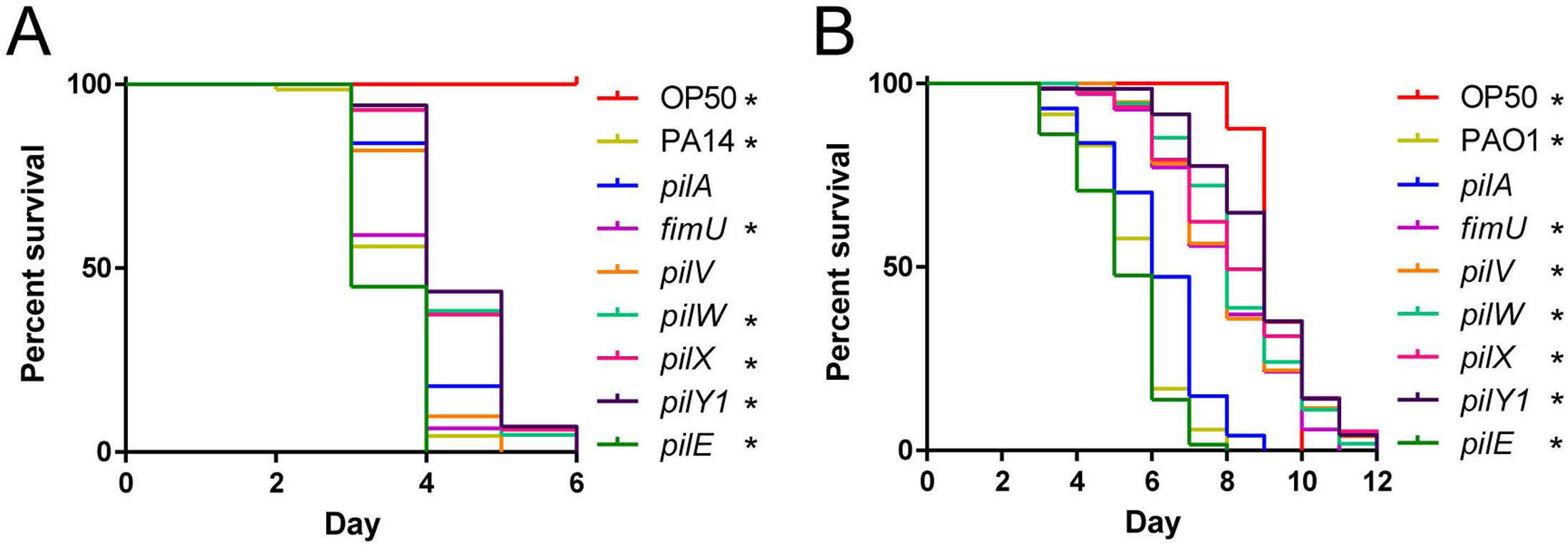
PilWXY1 contribute to T4P-independent virulence. (A) SK assays for PA14 *pilA*, *fimU*, *pilV*, *pilW*, *pilX*, *pilY1*, and *pilE* mutants. Synchronized L4 worms were seeded onto SK plates and scored for death every 24 h, then plotted as “percent survival” over the course of the assay. “Day” represents the number of days after L4 on which the plates were scored. PA14 *fimU* and *pilE* mutants had similar virulence to WT, *pilA* and *pilV* mutants were slightly less virulent than WT, and *pilW, pilX,* and *pilY1* mutants were less virulent than all other strains tested. (B) SK assays for PAO1 *pilA*, *fimU*, *pilV*, *pilW*, *pilX*, *pilY1*, and *pilE* mutants. The PAO1 *pilE* mutant had similar virulence to WT, the *pilA* mutant was slightly less virulent, and *fimU*, *pilV*, *pilW*, *pilX*, and *pilY1* mutants were much less virulent. In (A) and (B), asterisks indicate strains that were significantly different from a *pilA* mutant by Gehan-Breslow-Wilcoxon test at p = 0.05 (p = 0.00625 with a Bonferroni correction), n = 3 trials.

### PilWXY1 promote virulence in a SadC-independent manner

PilWXY1 were previously proposed to increase c-di-GMP production by SadC, such that loss of *pilW*, *pilX*, or *pilY1* resulted in a biofilm-deficient phenotype, indicative of low intracellular c-di-GMP [39, 40, 43]. Therefore, we hypothesized that biofilm defects of *pilW*, *pilX*, and *pilY1* might impede their ability to colonize the *C. elegans* gut, leading to reduced virulence. The PA14 and PAO1 parent strains and their cognate *pilA*, *fimU*, *pilV*, *pilW*, *pilX*, *pilY1*, and *pilE* mutants formed negligible levels of biofilm in liquid SK medium, chosen to approximate the growth conditions used for the SK assay (Supplementary Fig S2). To assess the levels of cyclic-di-GMP in these strains, we constructed a luminescence-based *cdrA* promoter reporter based on an extensively-characterized green fluorescent protein-based reporter system [44, 49-54]. *cdrA* promoter activity has been positively correlated with c-di-GMP levels, as measured by liquid chromatography-mass spectrometry [49, 51, 53, 54]. We verified that overexpression of SadC led to a ~60-fold increase in *cdrA* promoter activity, while overexpression of AlgR, which positively regulates genes that promote c-di-GMP production [55, 56], led to a ~2-fold increase in promoter activity that was enhanced to ~4-fold when *algR* expression was increased with 0.05% L-arabinose (Fig 2A). Deletion of *sadC* or *algR* led to a ~2-fold decrease in *cdrA* promoter activity relative to WT. *cdrA* promoter activity in WT is expected to be relatively low in liquid media because c-di-GMP levels increase upon surface attachment [43]. Compared to WT, *pilW*, *pilX*, and *pilY1* had ~3-fold lower *cdrA* promoter activity, indicative of reduced c-di-GMP (Fig 2B). These results are consistent with reports that PilWXY1 promote c-di-GMP production via SadC [39, 40, 43]. We next investigated whether SadC was required for virulence towards *C. elegans*, as would be predicted if decreased virulence in *pilW*, *pilX*, and *pilY1* mutants was due to dysregulation of SadC activity. A small decrease in virulence towards *C. elegans* was previously reported for a PA14 *sadC* mutant [57]; however, we saw no difference in virulence between WT and a *sadC* mutant in either the PA14 or PAO1 backgrounds (Supplementary Fig S3). Further, overexpression of SadC led to a hyper-biofilm phenotype *in vitro* in SK medium, but a slight reduction in virulence, demonstrating that the amount of biofilm formed *in vitro* does not correlate with virulence in *C. elegans* (Fig 3). Although the exact mechanisms of *P. aeruginosa* pathogenesis in *C. elegans* are not fully understood, biofilms were suggested to be important for establishment of infection [57-59]. Our *in vitro* biofilm data suggests that biofilms may not be a major contributor to *P. aeruginosa* pathogenesis in this model, but direct visualization and quantification of biofilms within the nematode gut will be required to support this conclusion.

**Fig 2.**
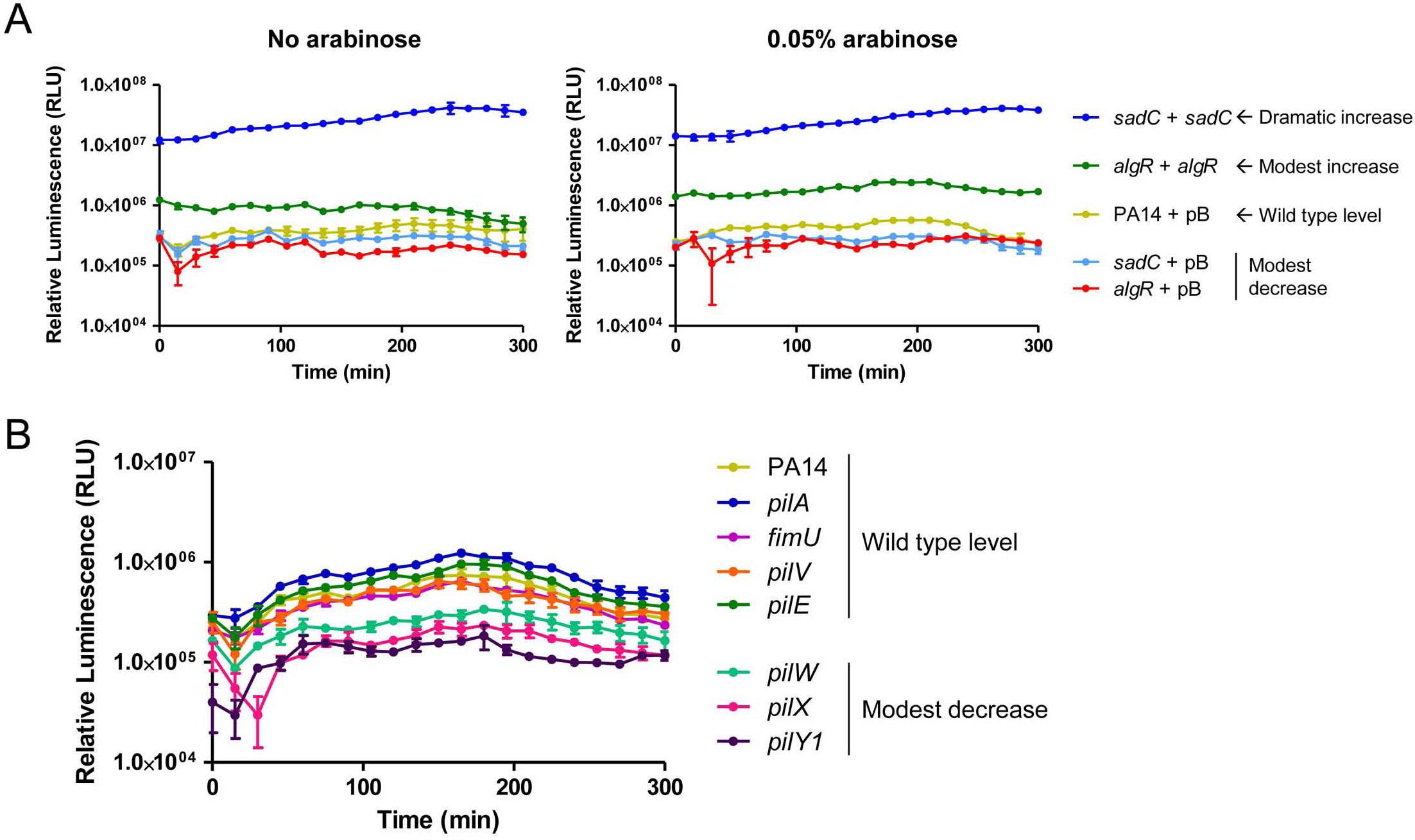
*pilW*, *pilX*, and *pilY1* mutants have reduced *cdrA* promoter activity. (A) *cdrA* promoter activity in PA14 *sadC* and *algR* deletion and overexpression strains. pMS402-P*cdrA*, containing the *lux* genes under expression of the *cdrA* promoter, was introduced into strains of interest, along with pBADGr (vector-only control), pBADGr-*sadC*, or pBADGr- *algR*. Assays were set up in technical triplicate in SK media, with or without 0.05% L-arabinose to induce expression of the pBADGr promoter, and measurements were taken every 15 min over 5 h. Loss of *sadC* or *algR* led to a subtle decrease in *cdrA* promoter activity, while SadC overexpression led to a dramatic increase in *cdrA* promoter activity. Overexpression of AlgR also led to a subtle increase in *cdrA* promoter activity that was enhanced upon addition of L-arabinose. n = 3 trials. (B) *cdrA* promoter activity in PA14 *pilA*, *fimU*, *pilV*, *pilW*, *pilX*, *pilY1*, and *pilE* mutants. Loss of *pilW*, *pilX*, or *pilY1* led to a decrease in *cdrA* promoter activity. n = 3 trials.

**Figure 3.**
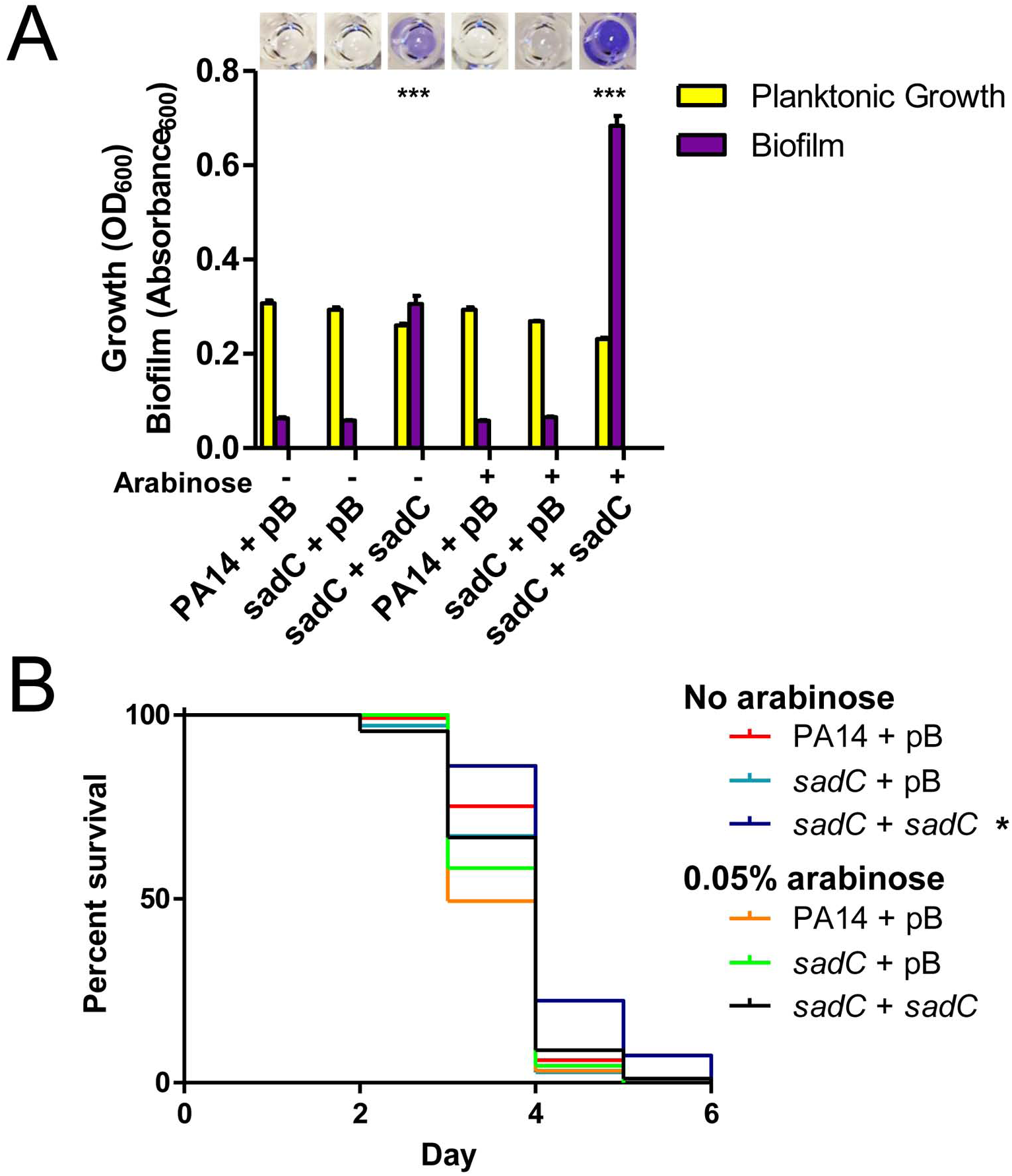
SadC promotes biofilm formation but is not required for virulence. (A) Biofilm assays for *sadC* deletion and overexpression strains. PA14 *sadC* biofilm levels were similar to WT. Expression of SadC *in trans* from a multicopy plasmid led to increased biofilm formation relative to WT at 0% (due to leaky promoter) and 0.05% L-arabinose, p < 0.001. Significance was determined by one-way ANOVA followed by Dunnett post-test relative to PA14 + pBADGr, n = 3. (B) SK assays for *sadC* deletion and overexpression strains. Overexpression of SadC led to a subtle but reproducible loss of virulence relative to WT at 0% L-arabinose. A *sadC* mutant had WT virulence. Asterisks indicate strains that were significantly different from PA14 + pBADGr by Gehan-Breslow-Wilcoxon test at p = 0.05 (p = 0.0125 with a Bonferroni correction), n = 3.

### PilVWXY1 repress expression of the MP operon

After ruling out involvement of the SadC pathway, we next explored the potential role of FimS-AlgR in PilWXY1-mediated virulence. Informed by previous work in our laboratory showing that the sensor kinase PilS of the PilSR TCS interacts directly with PilA in the inner membrane to decrease PilR-dependent major pilin expression [60], we hypothesized that FimS interacts with one or more MPs, and that loss of that interaction could lead to activation of AlgR and subsequent upregulation of the MP operon. Bacterial two-hybrid (BACTH) assays were used to identify potential interactions between FimS and PilA, FimU, PilV, PilW, PilX, or PilE (Fig 4A). We also screened for interaction of FimS and AlgR, which has been inferred but never demonstrated [19]. Interactions between FimS and each pilin were identified; however, based on our experience with PilS [60], binding of pilins is necessary but not sufficient for inhibition. We also demonstrated interaction of FimS and AlgR (Fig 4A), providing further support for the hypothesis that FimS is the sensor kinase for AlgR.

**Fig 4.**
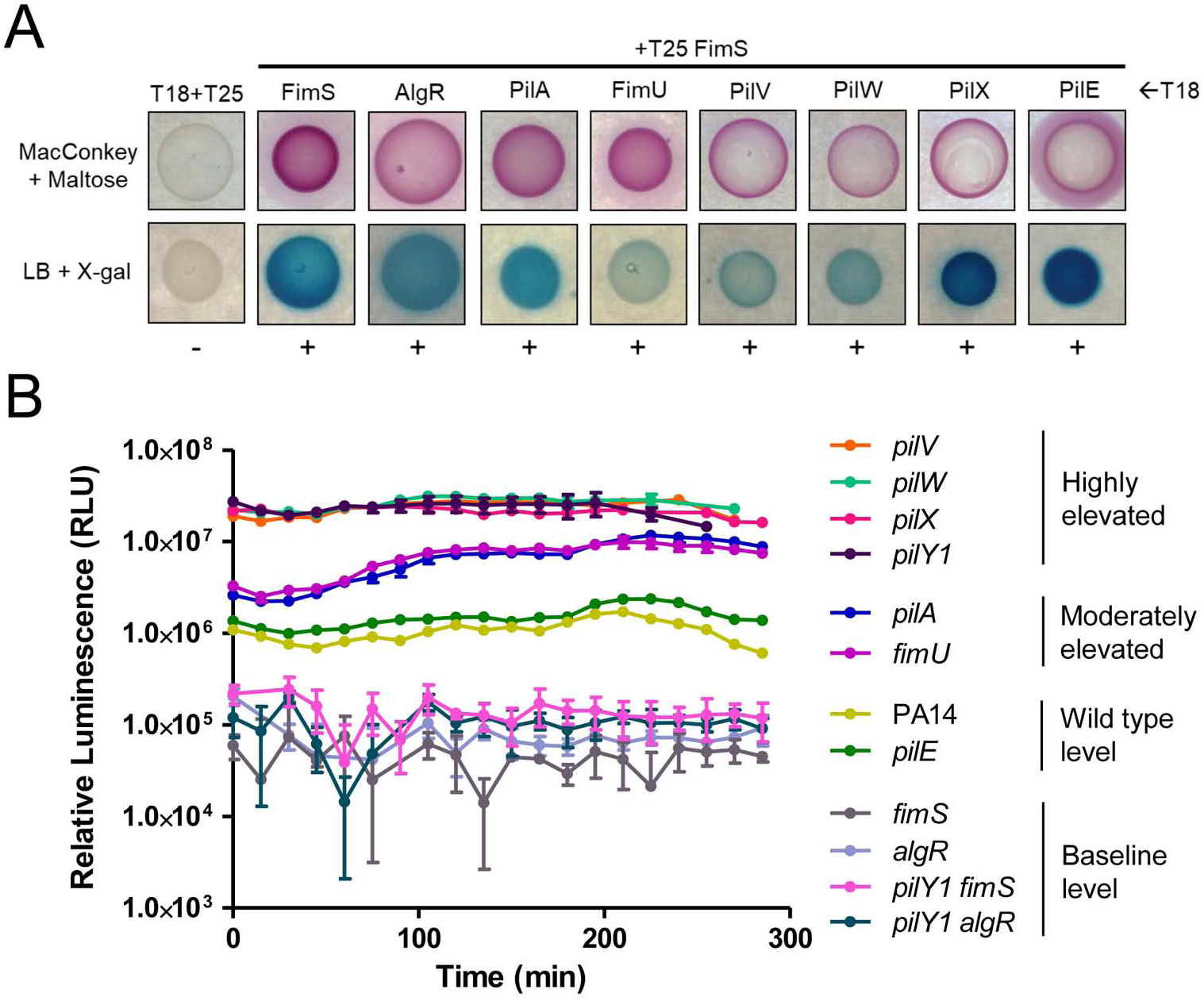
PilVWXY1 repress their expression via FimS-AlgR. (A) BACTH assays for FimS, AlgR, PilA, and MPs. Protein fusions with T18 and T25 fragments of the CyaA adenylate cyclase were screened for interactions on MacConkey and LB + X-gal plates. FimS interacted with AlgR, PilA, FimU, PilV, PilW, PilX, and PilE. Positive (+) or negative (-) interactions are indicated below each image, n = 3. (B) *fimU* promoter activity in PA14 *pilA*, *fimU*, *pilV*, *pilW*, *pilX*, *pilY1*, *pilE*, *fimS*, *algR*, *pilY1 fimS*, or *pilY1 algR* mutants. pMS402-P*fimU*, containing the *fimU* promoter upstream of the *lux* genes, was introduced into strains of interest. Loss of *pilV*, *pilW*, *pilX*, or *pilY1* led to highly elevated *fimU* promoter activity. *pilA* and *fimU* mutants had moderately increased *fimU* promoter activity relative to WT. *fimS* and *algR* mutants had negligible luminescence, and loss of *fimS* or *algR* also reverted *fimU* promoter activity in the *pilY1* mutant to baseline. n = 3 trials.

To decipher which MPs might modulate expression of the operon, we monitored expression from the *fimU* promoter using a *luxCDABE* reporter. Compared to WT PA14, there was a ~25-fold increase in luminescence in *pilV*, *pilW*, *pilX*, and *pilY1* mutants, which could be restored to WT levels by expressing the corresponding pilin *in trans* (Fig 4B, Supplementary Fig S4). *fimU* and *pilA* mutants had ~5-fold increased promoter activity, while a *pilE* mutant was comparable to WT. *fimS* and *algR* mutants had low baseline luminescence, ~10-fold lower than WT. To determine whether the increased promoter activity in *pilV*, *pilW*, *pilX*, and *pilY1* mutants depended on FimS-AlgR, either *fimS* or *algR* was deleted in the *pilY1* mutant background. The *pilY1 algR* double mutant had low luminescence (~10-fold lower than WT), consistent with AlgR acting as a positive regulator of the MP operon [40]. Loss of *fimS* in the *pilY1* mutant background also abolished *fimU* promoter activity (~10-fold lower than WT), supporting the idea that FimS may monitor PilVWXY1 levels and activate AlgR when levels are low. Based on these data, PilA, FimU, and PilE are unlikely to modulate FimS-AlgR activity even though they can interact with FimS.

PilVWXY1 were previously proposed to form a complex in the inner membrane, such that loss of any one component destabilizes the others [24]. Since PilY1 is thought to be cleaved on the periplasmic side of the inner membrane, it is unlikely to interact directly with the transmembrane domains of FimS [24]. Thus, we suspected that high *fimU* promoter activity in the *pilY1* mutant was due to reduced levels of one or more of the other pilins. To address this, we overexpressed FimU, PilV, PilW, PilX, or PilE in the *pilY1* mutant and measured *fimU* promoter activity. All these strains had luminescence comparable to the *pilY1* mutant (Supplementary Fig S4). Conversely, distinct effects have been observed in other studies upon overexpression of PilY1 [39, 40, 43]. Therefore, we overexpressed PilY1 in the *pilW* and *pilX* (high-luminescence) backgrounds; but PilY1 alone was insufficient to alter *fimU* promoter activity. Together, the data suggest that no individual component of the PilVWXY1 subcomplex is capable of modulating FimS activity when others are absent.

We also tested whether PilD processing of PilVWX was required for modulation of FimS activity. We constructed a *pilD* mutant, which lacks twitching motility since unprocessed pilins remain in the inner membrane [23, 61]. The absence of *pilD* had no impact on *fimU* promoter activity (Supplementary Fig S5), and a *pilD* mutant had virulence equivalent to a *pilA* mutant, likely attributable to its lack of T4P. Thus, PilVWX can modulate FimS activity in their unprocessed form.

### Hyperactivation of AlgR attenuates virulence

Because the results suggested that loss of PilWXY1 relieves feedback inhibition on FimS-AlgR, resulting in AlgR activation, we tested whether hyperactivation of AlgR alone could decrease virulence. We made chromosomal *algR*_D54E_ phospho-mimetic point mutants [62] in both PA14 and PAO1 backgrounds. We also made *algR*_D54A_ point mutants, as AlgR phosphorylation is required for transcription of a subset of genes in its regulon, including the MP operon [17, 62, 63]. We verified that the *algR*_D54A_ mutant was defective for twitching motility, while the *algR*_D54E_ mutant had WT twitching (Supplementary Fig S6). Unexpectedly, a *fimS* mutant retained ~50% twitching motility, in contrast to previous reports [18, 62]. In the absence of FimS, AlgR might be phosphorylated by small phosphate donors [64]. Based on the *fimS* data, we also questioned the assumption that AlgR phosphorylation was necessary for expression from the *fimU* promoter. When we overexpressed WT AlgR or AlgR_D54A_ in the *algR* mutant (Supplementary Fig S6), its twitching defect was fully complemented by AlgR, and partially complemented (25%) by AlgR_D54A_. Thus, although it increases binding to the *fimU* promoter [17, 62], phosphorylation of AlgR is not essential for transcription of the MP operon.

SK assays were then performed for PA14 and PAO1 *algR*_D54A_ and *algR*_D54E_ mutants, plus PA14 and PAO1 *fimS* and *algR* deletion mutants. PA14 and PAO1 *algR*_D54E_ mutants were less pathogenic than the corresponding WT strains, while *fimS*, *algR* and *algR*_D54A_ mutants had WT virulence (Fig 5AB). Loss of FimS-AlgR decreases expression of the MPs and PilY1 and prevents pilus assembly [17, 40]. Because our data show that loss of FimS-AlgR (and thus MP expression) had no impact on virulence, we conclude that reduced virulence of *pilW*, *pilX*, and *pilY1* mutants is due to the resulting activation of FimS-AlgR.

**Fig 5.**
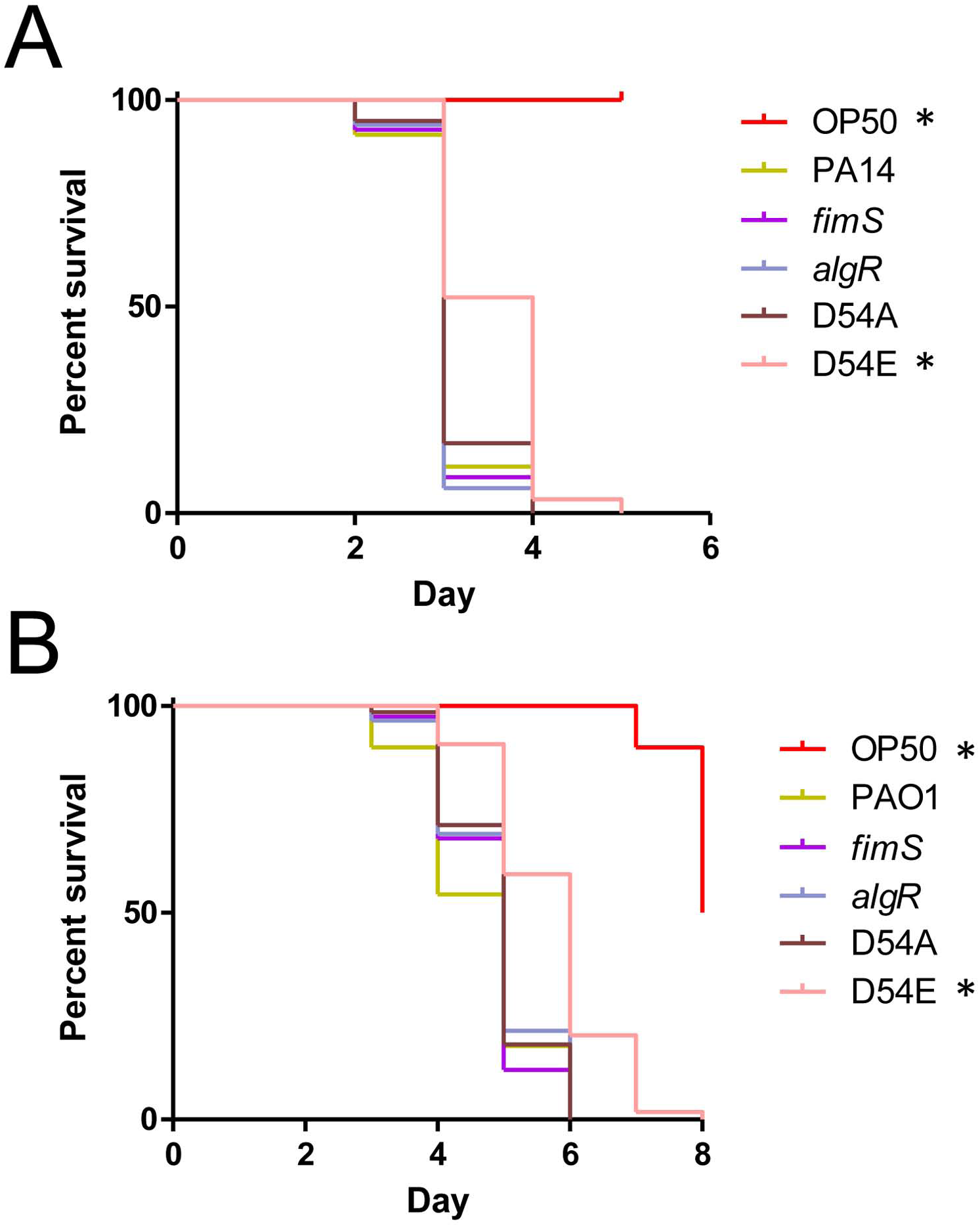
AlgR hyperactivation reduces virulence. SK assays for (A) PA14 and (B) PAO1 *fimS*, *algR*, *algR*_D54A_, and *algR*_D54E_ mutants. The *fimS*, *algR,* and *algR*_D54A_ mutants had WT virulence, while the *algR*_D54E_ mutants were less virulent than WT. For (A) and (B), asterisks indicate strains that were significantly different from WT by Gehan-Breslow-Wilcoxon test at p = 0.05 (p = 0.01 with a Bonferroni correction), n = 3 trials.

### Overexpression of AlgR attenuates virulence

Increased transcription of *fimS*-*algR* in a *pilY1* mutant relative to WT has been reported [38], suggesting that reduced virulence could arise through expression of increased amounts of the FimS-AlgR TCS, as well as its activation. Therefore, we asked whether increased AlgR levels would attenuate virulence, as previously demonstrated in a mouse infection model [65]. When *algR* was expressed *in trans* from a multicopy plasmid in PA14 *algR*, virulence was reduced compared to the vector control (Fig 6A). Because un-phosphorylated AlgR can also affect transcription of a subset of genes [66, 67], we tested the same mutant complemented with AlgR_D54A_. Complementation of the *algR* mutant with AlgR_D54A_ resulted in a severe virulence defect relative to the vector-only control. Thus, AlgR hyperactivation and overexpression independently diminish *P. aeruginosa* virulence towards *C. elegans*. Lastly, as AlgR is a positive regulator of biofilm formation [17, 55, 56], we performed biofilm assays for PA14 *algR* complemented with AlgR or AlgR_D54A_. Expression of either variant led to hyper-biofilm formation (Fig 6B), further emphasizing that the ability of a strain to form biofilms in SK medium does not correlate with virulence in worms. Instead, we suggest that virulence factors repressed by FimS-AlgR are important for *C. elegans* SK, and an increase in AlgR levels and/or activity attenuates virulence.

**Fig 6.**
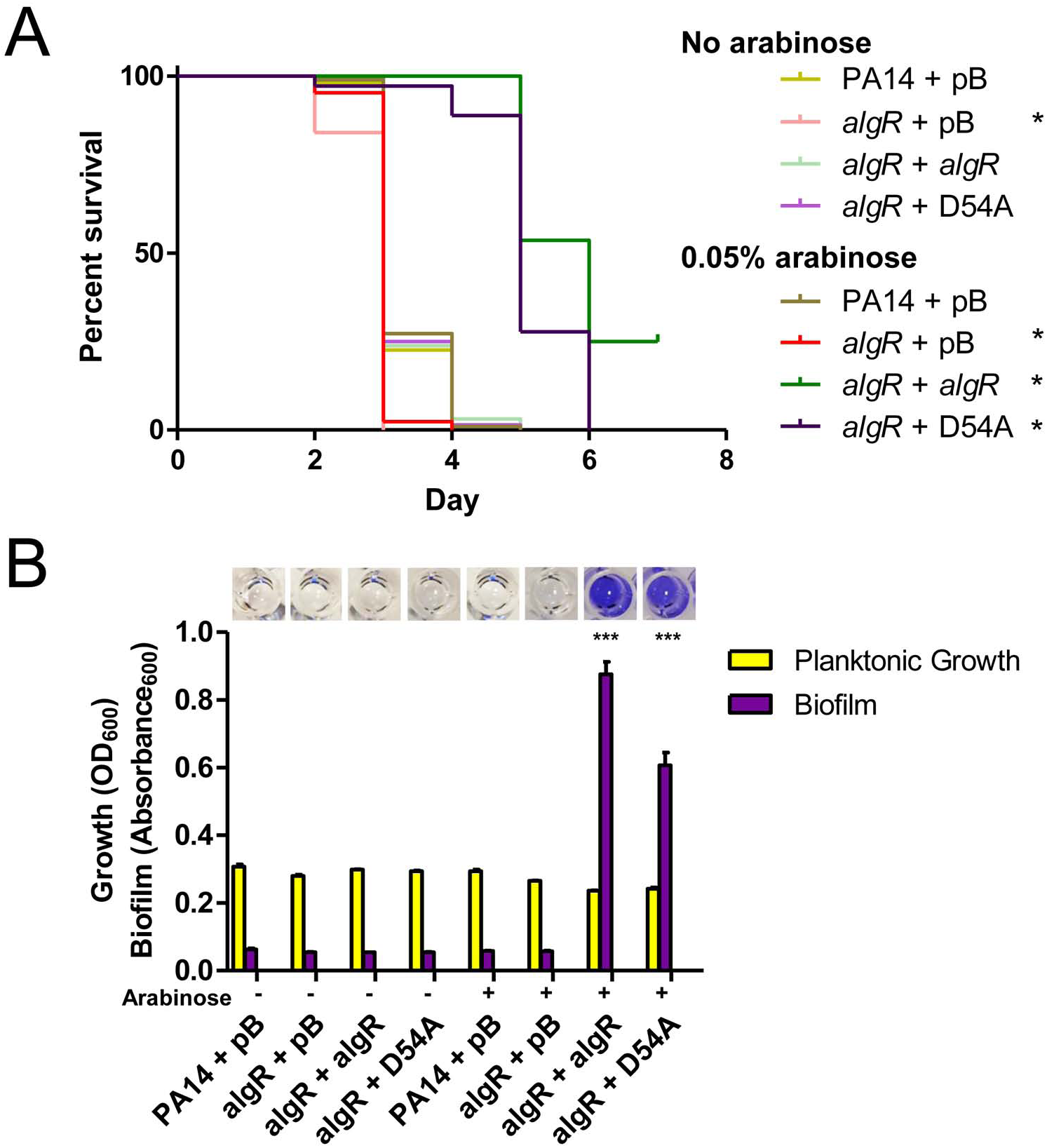
AlgR promotes biofilm formation and represses virulence. (A) SK assays for *algR* deletion and overexpression strains. Loss of *algR* led to a small increase in virulence, while overexpression of pBADGr-*algR* or pBADGr-*algR*_D54A_ reduced virulence at 0.05% L-arabinose. Asterisks indicate strains that were significantly different from PA14 + pBADGr by Gehan-Breslow-Wilcoxon test at p = 0.05 (p = 0.00833 with a Bonferroni correction), n = 3 trials. (B) Biofilm assays for *algR* deletion and overexpression strains. Microtiter plate biofilm assays were performed in liquid SK media over 24 h, in triplicate. Biofilms were stained with 1% crystal violet then solubilized in acetic acid. Loss of *algR* had no effect on biofilm formation. When grown at 0.05% L-arabinose, overexpression of pBADGr-*algR* or pBADGr-*algR*_D54A_ increased biofilm formation, p < 0.001. Significance was determined by one-way ANOVA followed by Dunnett post-test relative to WT, n = 3 trials.

### The virulence defects of *pilW*, *pilX* and *pilY1* mutants are dependent on FimS-AlgR

To provide further support for this model, we asked whether the virulence defects of PA14 *pilW*, *pilX*, and *pilY1* mutants required FimS-AlgR. We deleted *fimS* or *algR* in the *pilW*, *pilX*, and *pilY1* backgrounds, and tested virulence of the double mutants (Fig 7). We also deleted *pilW*, *pilX*, and *pilY1* in the *algR*_D54A_ background, to test if AlgR activation was necessary for the loss of virulence in *pilW*, *pilX*, and *pilY1* mutants. In all cases, the double mutants had WT virulence, equivalent to that of the *fimS*, *algR*, or *algR*_D54A_ single mutants. These results demonstrate that decreased virulence resulting from loss of PilWXY1 requires both FimS and AlgR. Although overexpression of AlgR_D54A_ *in trans* repressed virulence (Fig 6A), the chromosomal mutation was sufficient to alleviate the virulence defect of *pilW*, *pilX*, and *pilY1* mutants, suggesting that AlgR phosphorylation is important for PilWXY1-modulated virulence.

**Fig 7.**
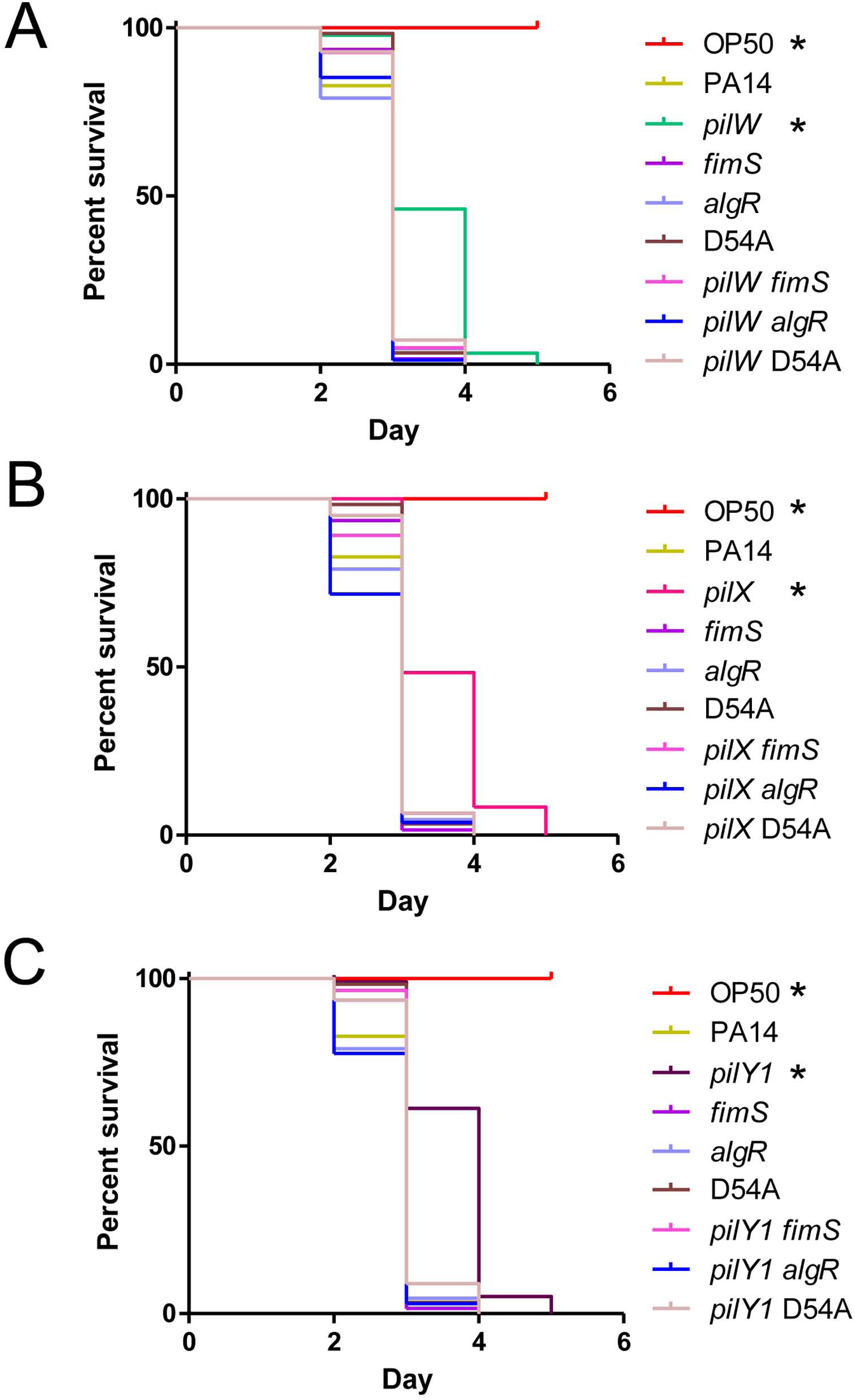
The virulence defect of *pilW*, *pilX*, and *pilY1* mutants is dependent on FimS-AlgR. SK assays for *pilW*, *pilX*, *pilY1*, *fimS*, *algR*, and *algR*_D54A_ single and double mutants. *fimS*, *algR*, and *algR*_D54A_ mutants have WT virulence. *pilW*, *pilX*, and *pilY1* have reduced virulence relative to WT, *fimS*, *algR*, and *algR*_D54A_ mutants. Combination of *pilW*, *pilX*, or *pilY1* mutations with *fimS*, *algR*, or *algR*_D54A_ mutations results in virulence equivalent to *fimS*, *algR*, and *algR*_D54A_ single mutants, respectively. All graphs represent 1 trial, separated into 3 graphs where strains relevant to (A) *pilW*, (B) *pilX*, and (C) *pilY1* mutants are included. Asterisks indicate strains that were less virulent than PA14 by Gehan-Breslow-Wilcoxon test at p = 0.05 (p = 0.003125 with a Bonferroni correction), n = 3.

The sigma factor AlgU (AlgT/σ^22^/σ^E^) acts upstream of FimS-AlgR to promote *algR* transcription [68-70], thus we tested its potential involvement in modulation of virulence by PilWXY1. An *algU* mutant was more virulent than WT (Fig 8), as previously demonstrated in mouse models [71], while *pilW algU*, *pilX algU*, and *pilY1 algU* double mutants had near-WT virulence (less than an *algU* mutant, but more than *pilW*, *pilX*, and *pilY1* single mutants).

**Fig 8.**
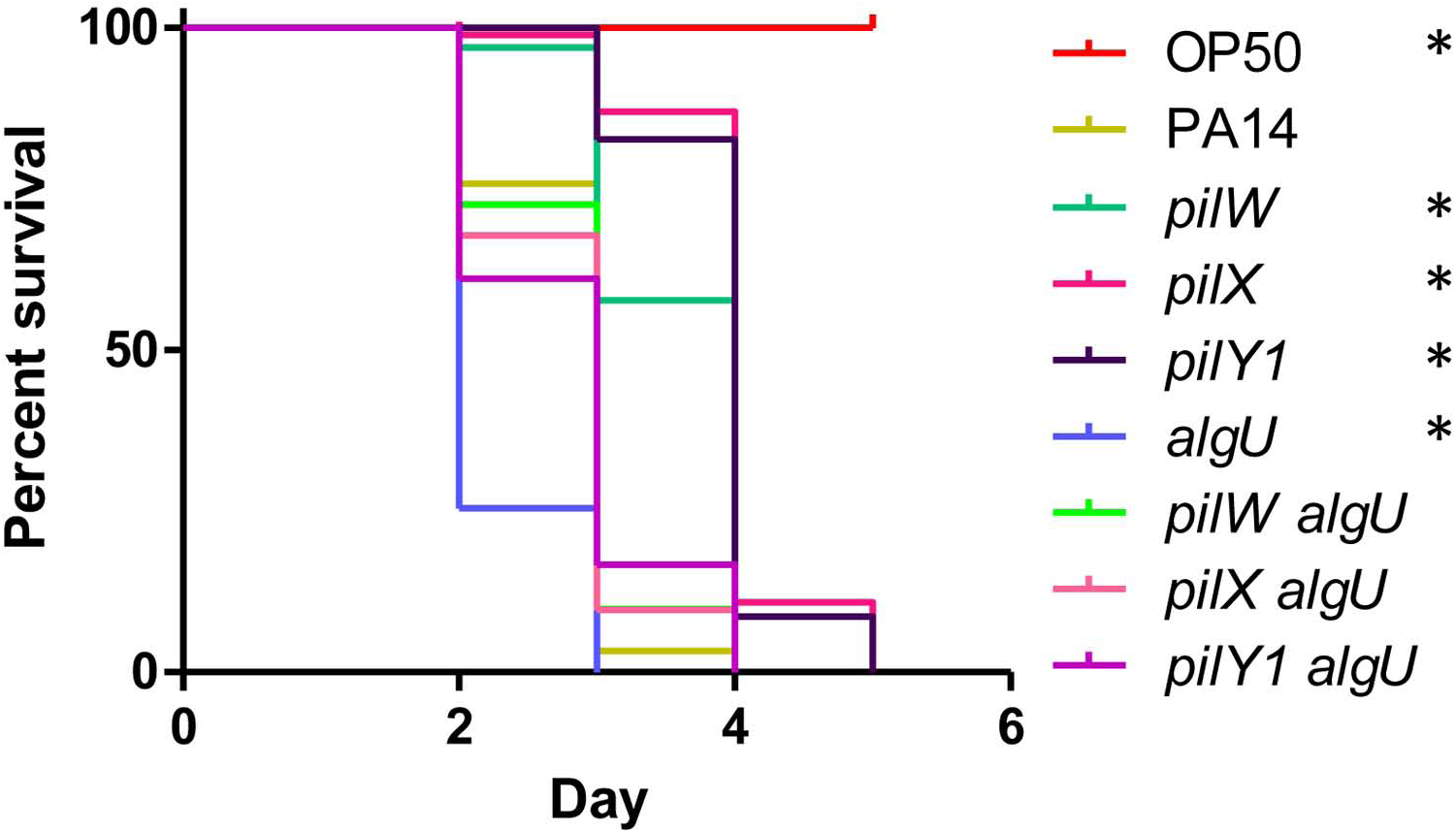
PilWXY1-mediated virulence is not dependent on AlgU. SK assays for PA14 *pilW*, *pilX*, *pilY1*, *algU*, *pilW algU*, *pilX algU*, and *pilY1 algU* mutants. Loss of *algU* led to increased pathogenicity relative to WT, while *pilW algU*, *pilX algU*, and *pilY1 algU* mutants had near-WT virulence. Asterisks indicate strains that were significantly different from PA14 by Gehan-Breslow-Wilcoxon test at p = 0.05 (p = 0.00625 with a Bonferroni correction), n = 3.

Although AlgU promotes *algR* transcription [69], loss of AlgU alone does not prevent expression of AlgR [68]. Given the reduced virulence of the *pilW algU*, *pilX algU*, and *pilY1 algU* double mutants relative to *algU*, PilWXY1 modulation of FimS-AlgR signalling appears to be intact in the *algU* mutant. These data are consistent with studies showing that *mucA* and *mucD* mutants, in which *algR* and *algU* are highly transcribed [69, 72-74], are less virulent towards *C. elegans* [75-77].

## Discussion

*P. aeruginosa* uses T4P to attach to surfaces and host cells, for biofilm maturation, and to move across surfaces via twitching motility [2]. The MPs and PilY1 are important players in T4P biogenesis and function, but also in regulation of swarming motility, surface attachment, mechanosensation, and virulence [38-40, 43]. The MP operon is positively regulated by FimS-AlgR, a TCS implicated in regulation of chronic *P. aeruginosa* lung infections [17-19]. Here, we explored the connection between loss of PilWXY1 (and thus, loss of T4P) and AlgR activation in virulence towards *C. elegans*, as summarized in Fig 9. We showed that *pilW*, *pilX*, and *pilY1* mutants were less virulent than WT or a *pilA* mutant, supporting the idea that PilWXY1 modulate virulence independently of their role in T4P assembly. We confirmed previous reports [23, 33, 40] that in the absence of *pilV*, *pilW*, *pilX*, or *pilY1*, expression of the MP operon is significantly increased, and that this requires FimS-AlgR. Either hyperactivation or overexpression of AlgR reduced virulence, while loss of *fimS* or *algR* in *pilW*, *pilX*, or *pilY1* reverted virulence to WT levels.

**Fig 9.**
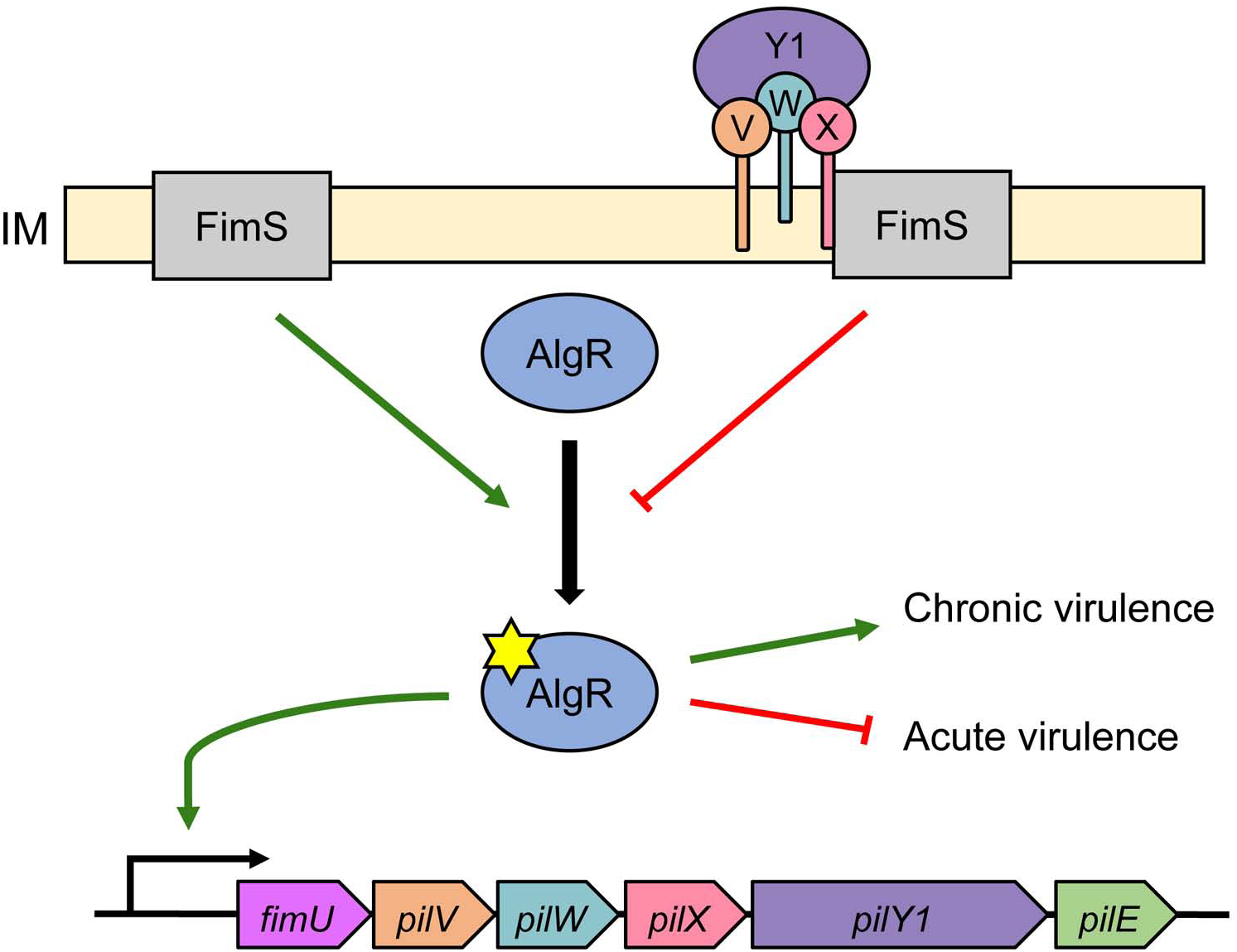
Hypothesized model for regulation of the MP operon via FimS-AlgR. When PilVWXY1 are absent, FimS may directly or indirectly promote increased protein levels and/or phosphorylation of AlgR (phosphate indicated by yellow star). Phospho-AlgR binds the *fimU* promoter to promote expression of *pilY1* and the MP genes. Phospho-AlgR also promotes expression of genes associated with chronic infections, and represses those associated with acute infections. As PilVWXY1 accumulate in the IM, they are likely detected by FimS, potentially leading to reduced AlgR protein levels and/or phosphorylation. Abbreviations: PilV, V (orange); PilW, W (cyan); PilX, X (pink); PilY1, Y1 (dark purple); IM, inner membrane.

These data – coupled with BACTH data showing that the minor pilins interact directly with FimS in the membrane (Fig 4) – suggest that FimS may act as a molecular thermostat to monitor MP levels, and in their absence, activates AlgR to upregulate expression of the MP operon. A similar inventory control mechanism was recently described for the PilSR TCS, where PilS phosphorylates PilR when PilA levels are low, and dephosphorylates PilR when PilA levels are high [60]. It is not yet clear if FimS responds to changes in levels of the PilVWXY1 subcomplex, thought to prime assembly of T4P [24, 78, 79]. When overexpressed individually *in trans*, each of the MPs inhibited twitching motility in PAO1 [23], but since the others were still expressed from the chromosome, the exact nature of the signal detected by FimS remains to be determined. When expressed *in trans*, no single component of the PilVWXY1 subcomplex reduced *fimU* promoter activity if others were absent (Supplementary Fig S4). The specific signal that inhibits FimS activity remains to be deciphered. Whether the FimS-inhibitory signal is the same in PA14 and PAO1 also remains unknown. Though PilWXY1 were required for virulence in PA14 and PAO1, FimU and PilV influenced virulence only in PAO1 (Fig 1A-B). Given the MPs are divergent, FimU and PilV may play different roles in PAO1 versus PA14 [48]. It is possible that FimU and PilV are more important for stability of the PilWXY1 subcomplex in PAO1 than in PA14, and/or that PAO1 FimU and PilV can directly modulate FimS activity.

Kuchma et al. [39, 43] reported that loss of *pilW*, *pilX*, or *pilY1* increased swarming motility and decreased biofilm formation, both indicative of low c-di-GMP levels. As biofilms were proposed to contribute to *P. aeruginosa* pathogenesis in *C. elegans*, we investigated whether the reduction in virulence in the absence of PilWXY1 was linked to decreased biofilm via loss of SadC activation [57-59, 80]. In our hands, levels of *sadC* had no impact on virulence even though they clearly modulated the amount of biofilm produced in SK media (Fig 3A-B, Supplementary Fig S3). Irazoqui et al. [59] examined the *C. elegans* gut during *P. aeruginosa* infection and described extracellular material that they suggested might indicate presence of a biofilm. Anti-biofilm compounds reduced *P. aeruginosa* virulence towards *C. elegans*, but a mechanism of action for those compounds has not been described [58]. Recently, the small RNA SrbA was shown to modulate both biofilm and virulence towards *C. elegans*; however, deletion of *srbA* led to altered transcription of at least 26 other genes that may also affect virulence [81].

Rather than using standard biofilm media, we performed these assays in liquid SK media to more closely mimic the conditions to which bacteria are exposed in the SK assay. To our knowledge, this is the first report to use SK media for biofilm assays. As we found no correlation between biofilm formation and virulence, we suggest that acute-phase virulence factors may be more important for *C. elegans* pathogenesis in the SK model. However, we recognize that *in vitro* biofilm assays may not replicate the conditions within the *C. elegans* gut; direct visualization of bacteria in worms will be needed to clarify the role of biofilm formation.

PilY1 and the MPs have been implicated in surface detection and activation of virulence, via signalling through SadC [38, 40]. Because loss of PilY1 or the MPs prevents T4P assembly and function, it is crucial to distinguish phenotypes resulting from lack of specific proteins versus loss of piliation [24]. Luo et al. [40] suggested that association of PilY1 with surfaces transduces a signal through the T4P machinery to stimulate c-di-GMP production by SadC, while Rodesney et al. [44] showed that loss of *pilA*, *pilY1*, or *pilT* prevents surface-activated c-di-GMP production. Rodesney et al. [44] proposed that both PilY1 and functional T4P are required for mechanosensation; however, it is not possible to delete *pilY1* without ablating T4P assembly. Our *cdrA* promoter reporter data support the idea that PilWXY1 promote cyclic-di-GMP production by SadC, as loss of *pilW*, *pilX*, or *pilY1* decreased *cdrA* promoter activity (Fig 2B). However, we argue that the PilWXY1-SadC pathway – though important for c-di-GMP signalling – is not critical for virulence towards *C. elegans*. Instead, our data show that PilWXY1-FimS-AlgR signalling axis is responsible for T4P-independent changes in virulence of *pilW*, *pilX*, and *pilY1* mutants. Thus, surface attachment may induce c-di-GMP production via PilWXY1-SadC [40, 43], while the brief trapping of T4P outside the cell upon contact with a surface might transiently deplete PilVWXY1 levels in the IM, resulting in increased FimS-AlgR activity and transition towards a sessile, biofilm lifestyle.

Whether the loss of *pilW*, *pilX*, or *pilY1* leads to increased amounts of AlgR, its increased phosphorylation via FimS, or both, remains to be clarified. Okkotsu et al. [62] showed that AlgR and AlgR_D54E_ levels are comparable, suggesting that the loss of virulence we observed for PA14 *algR*_D54E_ is attributable to the D54E phospho-mimetic mutation alone. Overexpression of AlgR_D54A_ *in trans* reduced virulence (Fig 6A), but the same mutation on the chromosome reverted virulence of *pilW*, *pilX*, and *pilY1* mutants to WT levels (Fig 7). Therefore, we suspect that it is primarily AlgR phosphorylation (or lack of AlgR dephosphorylation) that leads to decreased virulence. However, it is possible that both increased AlgR protein levels and phosphorylation contribute to the reduced pathogenicity of *pilW*, *pilX*, and *pilY1* mutants. Kong et al. [55] showed that AlgR binds *fimS*-*algR*, suggesting that the TCS could positively regulate its own transcription in response to reduced PilWXY1 levels.

In addition to being essential for T4P function, FimS and AlgR control alginate production in the context of chronic CF infections, where *algR* transcription is high [18, 82]. Phosphorylation of AlgR increases binding affinity at some – but not all – of its target sequences [17, 62, 63, 67]. For example, AlgR_D54N_ failed to support twitching motility, but did not affect alginate production [17, 63]. Our twitching motility data suggests that AlgR_D54A_ is capable of binding to the *fimU* promoter, albeit less efficiently than WT AlgR (Supplementary Fig S6). FimS is an unorthodox histidine kinase, with four transmembrane domains instead of the typical two, and lacks both a periplasmic sensing domain and the canonical motif involved in ATP coordination that mediates auto-phosphorylation [19, 83]. Direct interaction and/or phospho-transfer between FimS and AlgR have not been reported. Rather, the idea that FimS acts as a kinase for AlgR comes from this and other studies demonstrating similar phenotypes for *fimS, algR*, and *algR*_D54N_ mutants [17, 18, 84]. Here, we demonstrated that FimS and AlgR interact in the BACTH assay (Fig 4) lending further support to this model.

FimS and AlgR promote expression of genes important for production of alginate, biofilms, and c-di-GMP, and inhibit expression of virulence factors such as the T3SS, pyocyanin, and quorum sensing [55, 56, 74, 85, 86]. The observation that the loss of *algR* had no impact on virulence towards amoebae [38] or nematodes (Fig 5AB) suggests that the AlgR-activated genes may not contribute to virulence, although the mechanisms of killing could differ. In mouse models, *fimS* and *algR* deletion mutants are attenuated, though overexpression of AlgR also markedly reduces virulence [55, 65, 87]. Further, Little et al. demonstrated that PAO1 *algR*_D54E_ had WT virulence in *Drosophila melanogaster* and mouse infection models, while an *algR*_D54A_ mutant had highly attenuated virulence [87]. The outcomes that result from interaction of *P. aeruginosa* with different hosts will depend on a combination of factors including host defenses, site of infection, available nutrients, and virulence repertoire of a particular strain. However, our results suggest that changes in the specific repertoire of bacterial virulence factors, or the timing of their production, can tip the balance in the host’s favour.

The subset of AlgR-regulated virulence genes important for *C. elegans* pathogenesis is not defined. Screening of a PA14 transposon library for loss of virulence implicated several genes encoding regulators rather than individual virulence factors, suggesting that *C. elegans* pathogenesis is multifactorial [35]. Consistent with this hypothesis, a study of 18 WT *P. aeruginosa* strains revealed no correlation between pathogenicity and any specific virulence factors [88]. We saw WT or greater levels of virulence for *algR* and *algU* mutants, respectively, consistent with a role for AlgRU in repression of acute phase virulence factors (Figs 5, 8). Factors under positive control of AlgRU may be important during later stages of infection in more complex mammalian infection models, but not crucial for pathogenesis in nematodes [89, 90]. In support of this hypothesis, past studies have demonstrated that increased mucoidy, via mutation of *mucA* or *mucD*, reduced nematode killing [75-77].

While important for the initial stages of infection, T4P are less critical in chronic CF lung infections and are often lost over time [5, 91, 92]. *P. aeruginosa* CF isolates frequently become mucoid via activation of AlgR, and production of many virulence factors is reduced [82, 93, 94]. Although the two outcomes are not necessarily temporally or mechanistically linked, mutations that achieve both may be advantageous during chronic CF lung infections. Specifically, loss of PilWXY1 may be adaptive in the context of CF, leading to AlgR activation and loss of T4P function. To test this idea, it will be interesting to examine the genotypes of mucoid CF isolates for these types of mutations. In conclusion, our results suggest that PilWXY1 promote virulence towards *C. elegans* by inhibiting FimS-AlgR activation. These data demonstrate how loss of one virulence factor (T4P) may activate others (via AlgR). Because the interplay between virulence factors in *P. aeruginosa* is complex and dynamic, careful consideration will be required when designing potential anti-virulence therapeutic strategies.

## Materials and methods

### Bacterial strains and plasmids

Strains and plasmids used in this work are listed in Supplementary Table S1. Bacteria were grown at 37°C for 16 h in 5 ml lysogeny broth (LB) Lennox, or on 1.5% agar LB plates, unless otherwise specified. Plasmids were transformed into chemically-competent *E. coli* by heat-shock, and into *P. aeruginosa* by electroporation [95]. Where appropriate, gentamicin (Gm) was added at 15 µg/ml for *E. coli*, and 30 µg/ml for *P. aeruginosa*. Kanamycin (Kan) was added at 50 µg/ml for *E. coli*, and 150 µg/ml for *P. aeruginosa*. Ampicillin (Amp) was added at 100 µg/ml for *E. coli*. L-arabinose was added at 0.05% where indicated to induce expression from the pBADGr promoter [96].

### Cloning procedures

Vectors were constructed using standard cloning procedures, using the primers listed in Supplementary Table S2. Deletion constructs were designed to contain 500-1000 bp homology upstream and downstream the gene to be deleted. Deletion constructs for PA14 *fimU*, *pilV*, *pilW*, *pilX*, *pilY1*, and *pilE* were synthesized by Genscript in the pUC57Kan vector. pEX18Gm-*sadC* was created by amplifying the *sadC* deletion region from PA14 *sadC roeA* [42], followed by digestion and ligation into pEX18Gm. pEX18Gm-*fimS*, pEX18Gm-*algR*_D54A_, and pEX18Gm-*algR*_D54E_ were made by overlap extension PCR [97]. Restriction digestion followed by ligation of the upstream and downstream fragments was used to create the deletion constructs pEX18Gm-*algR*, pEX18Gm-*algU*, and pEX18Gm-*pilD*. pMS402-P*fimU* and pMS402-P*cdrA* were created by amplifying and digesting the promoter regions of the PA14 MP operon and *cdrA* gene, respectively. Digested pBADGr was treated with alkaline phosphatase prior to ligation to avoid re-circularization of the vector. Constructs were verified by Sanger sequencing (MOBIX lab, McMaster, Hamilton, ON).

### Mutant generation by allelic exchange

Allelic exchange was used to remove or alter specific genes [98]. pEX18Gm suicide plasmid derivatives (see Cloning procedures and Table 1) were used to create all mutants in this work. After heat-shock transformation into *E. coli* SM10 cells, pEX18Gm constructs were conjugated into corresponding PA14 or PAO1 parent strains. Cells were then transferred to *Pseudomonas* isolation agar (PIA) Gm100 plates and incubated for 18 h at 37°C, to select for integration of pEX18Gm derivatives into the chromosome. Colonies were streaked onto LB/sucrose and incubated at 30°C for 18 h to select against merodiploids. Resultant colonies were patched onto LB and LB Gm30 to identify gentamicin-sensitive colonies. Regions flanking the desired mutations were amplified and sequenced to confirm success.

### Twitching motility assays

Twitching motility assays were performed as previously described [99], with the following modifications. Individual colonies were stab-inoculated in triplicate into 1% agar LB solidified in plasma-treated tissue culture-grade plates (Thermo Fisher) and incubated at 30°C for 48 h. Agar was carefully removed and plates were stained with 1% crystal violet for 5 min.

Unbound dye was removed by rinsing with water, then stained twitching areas were measured using ImageJ. Twitching zones were normalized to WT (100%).

### Biofilm assays

Biofilm assays were performed as previously described, with modifications [100]. *P. aeruginosa* cultures were grown for 16 h at 37°C, diluted 1:200 in fresh LB, and grown to OD_600_ ~0.1. Cultures were then diluted 1:500 in liquid SK media (50 mM NaCl, 0.35% peptone, 1 mM CaCl_2_, 1 mM MgSO_4_, 5 µg/ml cholesterol in EtOH, 20 mM KH_2_PO_4_, and 5 mM K_2_HPO_4_), then 96-well plates were inoculated with 150 µl each strain, in triplicate. Sterility controls (liquid SK media) were included throughout the plate to check for contamination. Plates were covered with peg lids (Nunc) then wrapped in parafilm and incubated at 37°C for 24 h, shaken at 200 rpm. After incubation, the OD_600_ of the plate was measured to check for uniform growth and lack of contamination. Peg lids were washed for 10 min in 200 µl/well 1X phosphate-buffered saline (PBS), then stained with 200 µl/well 0.1% (w/v) crystal violet for 15 min. Unbound crystal violet was removed by washing lids in 70 ml distilled water 5 times at 10 min intervals. Crystal violet was solubilized from lids in 200 µl/well 33.3% acetic acid, then the absorbance at 600 nm was measured. Optical density and absorbance at 600 nm were plotted for growth and biofilm formation, respectively, then analyzed by one-way ANOVA followed by Dunnett post-test to compare each mutant to the WT control, p = 0.05. Error bars indicate standard error of the mean. Representative wells of acetic acid-solubilized crystal violet were imaged.

### *Caenorhabditis elegans* slow killing assay

SK assays were performed as described previously [101]. SK plates (0.35% peptone, 50 mM NaCl, 2% agar, 1 mM CaCl_2_, 5 µg/ml cholesterol, 1 mM MgSO_4_, 20 mM KH_2_PO_4_, 5 mM K_2_HPO_4_, 100 µM FUDR) were seeded with 100 µl of an overnight culture and incubated overnight at 37°C. The following day, plates were enriched with 1 ml of an overnight culture concentrated to 100 µl. Synchronized L4 worms were collected from *E. coli* OP50 plates, washed twice in M9 buffer, and then >50 worms were seeded onto each bacterial lawn on individual SK plates. SK plates were incubated at 25°C and scored for dead worms every 24 h. Worms were considered dead when they did not respond to touch, and were removed from the plate. OP50 was included as a negative control for virulence. Percent survival was plotted as a function of time. Survival curves were plotted on GraphPad Prism 5.00 for Windows, then compared using the Gehan-Breslow-Wilcoxon test, p = 0.05. Given that larvae were synchronized at 20°C then transferred at L4 to 25°C for the duration of the assay, worms were at risk of death due to senescence, rather than direct killing by *P. aeruginosa*, before day 10 [46]. Therefore, the Gehan-Breslow-Wilcoxon test, which gives weight to earlier timepoints, was used in favour of the standard log-rank test (notably, all reported differences were also significant by the standard log-rank test). To correct for multiple analyses, the critical p-value of 0.05 was divided by the number of pairwise comparisons made within an individual trial, as per the Bonferroni method [102]. Each assay was performed at least 3 times, and differences were only considered significant if they were reproducible in the majority of trials. Representative trials are shown; all replicates can be viewed in the Supplemental Material (Supplementary File S1).

### Luminescent reporter assay

Luminescent reporter assays were performed as previously described, with minor modifications [60]. Various strains harbouring the pMS402-P*fimU* or pMS402-P*cdrA* plasmids, encoding the luciferase genes under control of the *fimU* or *cdrA* promoters, respectively, were grown for 16 h at 37°C in LB Kan150, then diluted 1:50 in fresh liquid SK media with Kan150, in addition to Gm30 and 0.05% L-arabinose where appropriate. Subsequently, 100 µl of each culture was added to white-walled, clear-bottom 96-well plates (Corning) in triplicate, and incubated with shaking at 37°C in a Synergy 4 microtiter plate reader (BioTek). Luminescence readings were taken every 15 min for 5 h, and normalized to growth (OD_600_) at each time point. Readings that exceeded the limit of detection (>4 000 000 luminescence units) were discarded. At least 3 individual trials were performed. Error bars indicate standard error of the mean.

### Bacterial two-hybrid β-galactosidase activity assay

To test for interactions between FimS and AlgR or individual pilins, bacterial two-hybrid (BACTH) assays were performed as previously described [103]. pUT18C and pKT25 derivatives, encoding the T18 and T25 domains of the *Bordetella pertussis* CyaA adenylate cyclase fused to the N-terminus of FimS, AlgR, PilA, FimU, PilV, PilW, PilX, or PilE [24, 60, 104], were co-transformed into *E. coli* BTH 101 to screen for pairwise interactions. Single colonies were inoculated in 5 ml LB Amp100 Kan50 and grown overnight. The following day, 100 µl was inoculated into 5 ml fresh media and grown to OD_600_ = 0.6, then 5 µl was spotted onto MacConkey plates (1.5% agar, 100µg/ml ampicillin, 50µg/ml kanamycin, 1% (w/v) maltose, 0.5mM isopropyl b-D-thiogalactopyranoside) (Difco) or LB Amp100 Kan50 plates supplemented with 100 µl of 20 mg/ml X-gal. Plates were incubated at 30°C for 24 h. An interaction was considered positive when colonies appeared pink or blue on MacConkey and LB + X-gal plates, respectively. BTH 101 expressing pUT18C and pKT25 empty vectors was used as a negative control, and BTH 101 expressing pUT18C-*fimS* and pKT25-*fimS* was used as a positive control [49].

## Supporting information

**Fig S1.**
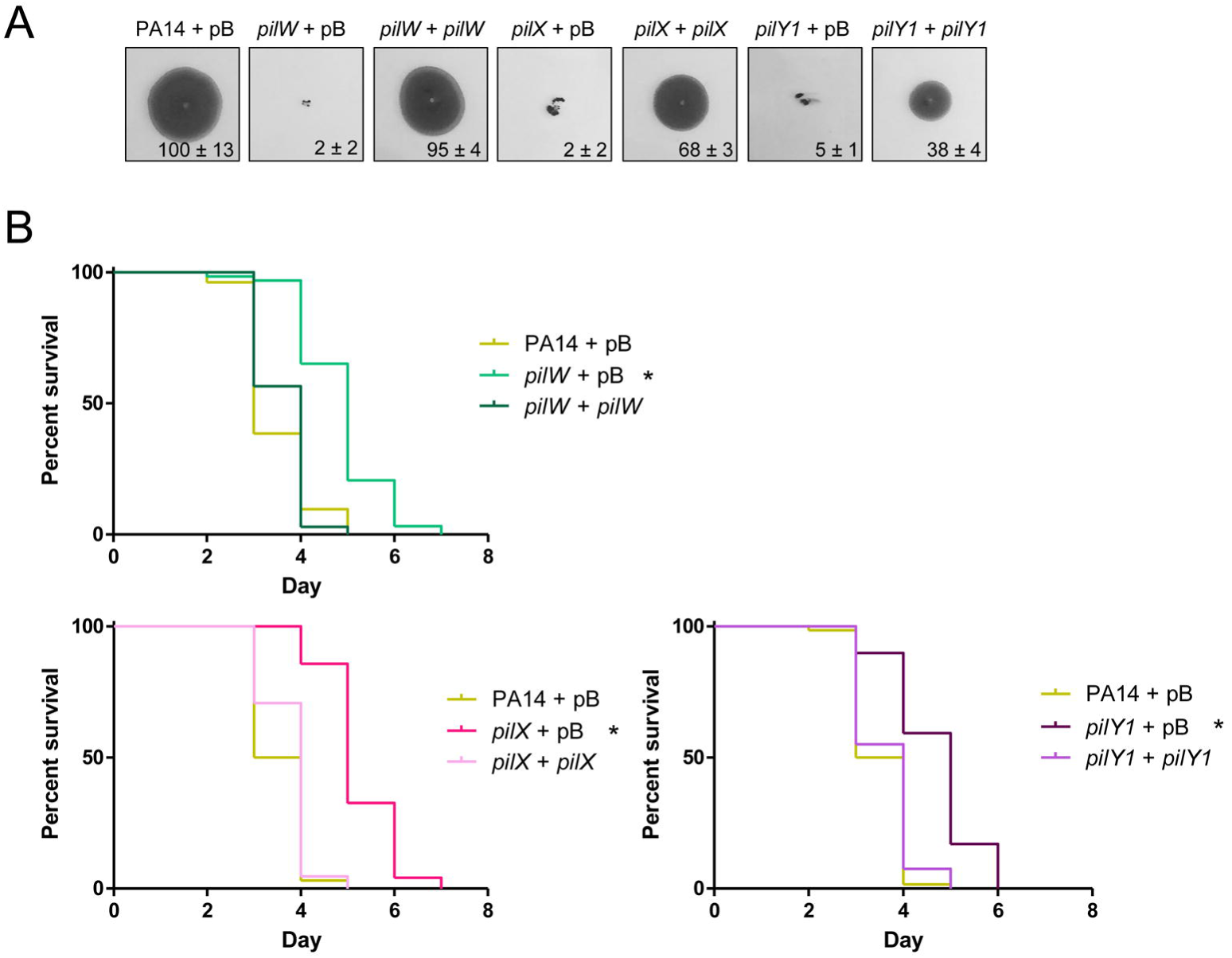
Twitching motility and virulence of *pilW*, *pilX*, and *pilY1* mutants can be complemented *in trans*. (A) Twitching motility assays for complemented PA14 *pilW*, *pilX*, and *pilY1* mutants. Colonies were stab-inoculated into 1% agar LB plates, in triplicate. Plates were stained with crystal violet after 48 h at 30°C. Complementation of PA14 *pilW*, *pilX*, and *pilY1* mutants with pBADGr-*pilW*, pBADGr-*pilX*, or pBADGr-*pilY1*, respectively, led to increased TM relative to complementation with pBADGr alone. Numbers indicate percent twitching area relative to WT, n = 3. (B) SK assays for complemented PA14 *pilW*, *pilX*, and *pilY1* mutants. Complementation of *pilW*, *pilX*, and *pilY1* mutants with pBADGr-*pilW*, pBADGr-*pilX*, or pBADGr-*pilY1*, respectively, restored virulence to near-WT levels. Asterisks indicate strains that were less virulent than PA14 + pBADGr by Gehan-Breslow-Wilcoxon test at p = 0.05 (p = 0.00833 with a Bonferroni correction), n = 3. Individual graphs represent separate trials.

**Fig S2.**
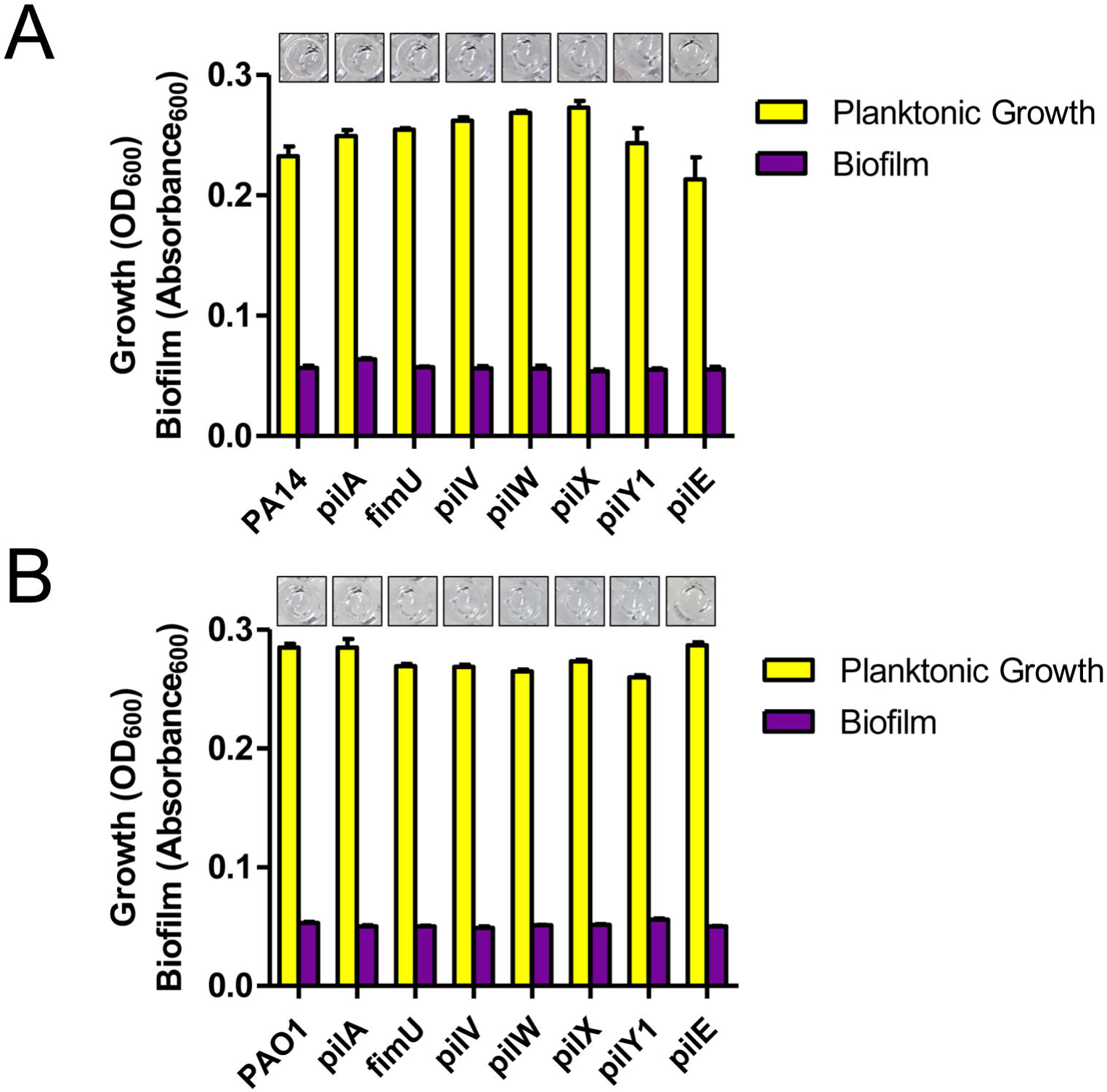
PA14 and PAO1 produce low levels of biofilm in liquid slow killing media. Biofilm assays for (A) PA14 and (B) PAO1 *pilA, fimU*, *pilV*, *pilW*, *pilX*, *pilY1*, and *pilE* mutants. Very little biofilm formation was detectable in liquid SK media for any strains. There were no differences in biofilm formation as determined by one-way ANOVA followed by Dunnett post-test relative to WT at p = 0.05, n = 3.

**Fig S3.**
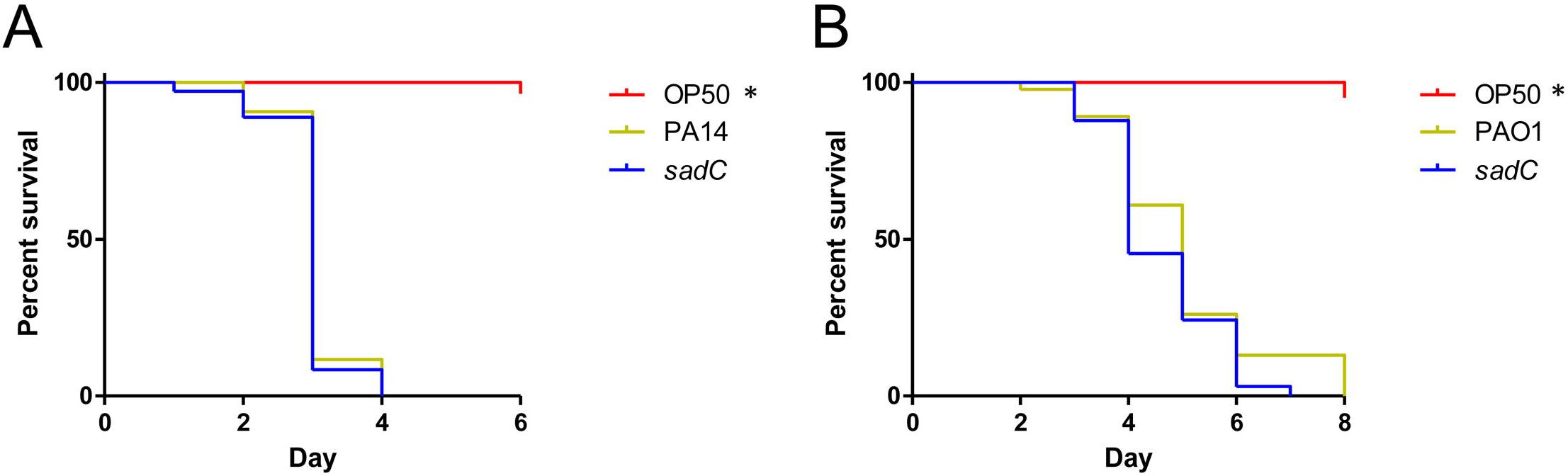
SadC is not required for virulence in PA14 or PAO1. SK assays for (A) PA14 and (B) PAO1 *sadC* mutants. Loss of *sadC* had no impact on pathogenicity relative to each respective WT strain, as measured by Gehan-Breslow-Wilcoxon test at p = 0.05 (p = 0.025 with a Bonferroni correction), n = 3.

**Fig S4.**
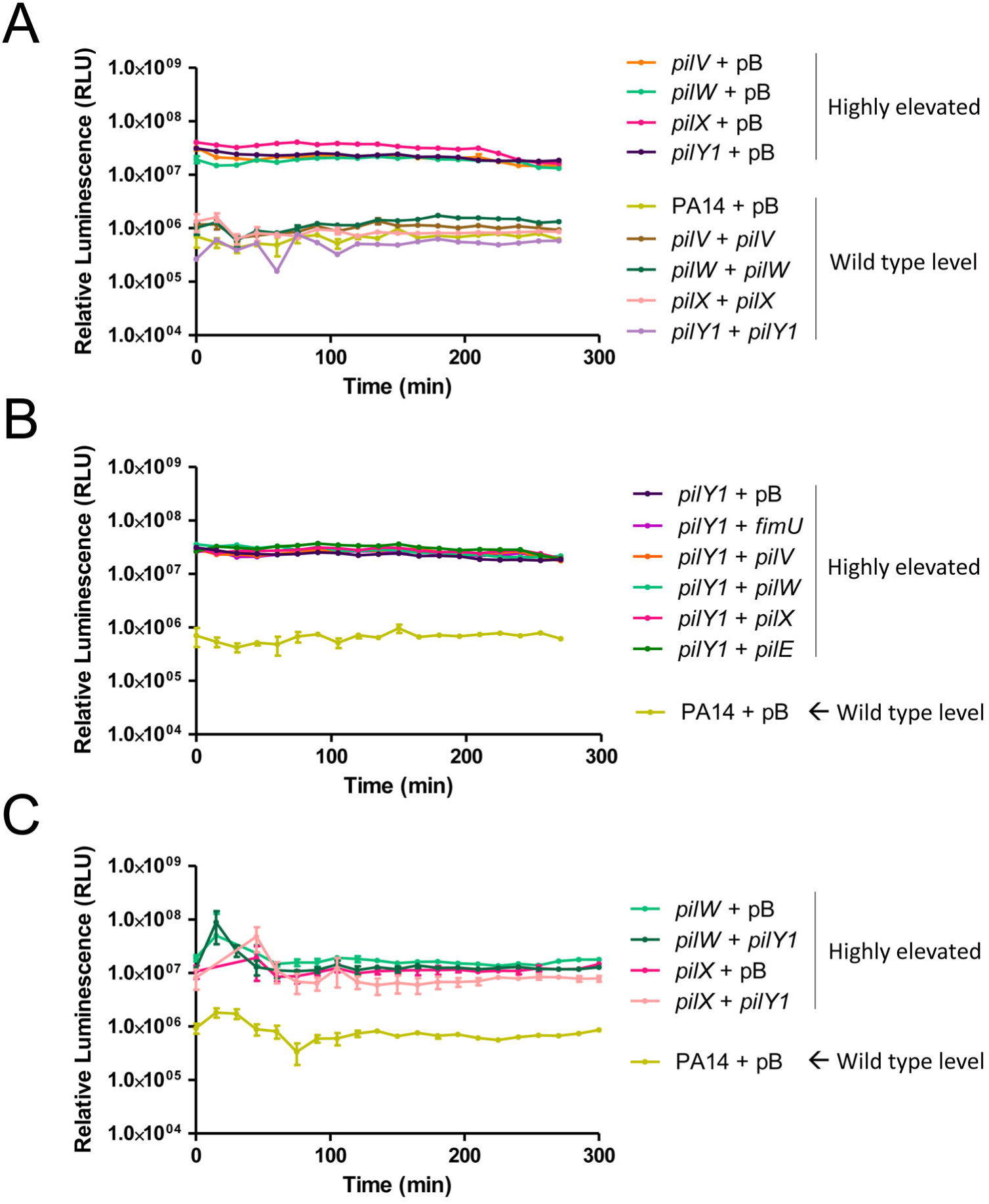
*pilV*, *pilW*, *pilX*, and *pilY1* mutants cannot be cross-complemented for *fimU* promoter activity. *fimU* promoter activity of *pilV*, *pilW*, *pilX*, and *pilY1* mutants complemented with the respective gene *in trans*. The high luminescence of each mutant was restored to WT level when *pilV*, *pilW*, *pilX*, and *pilY1* were complemented with PilV, PilW, PilX, and PilY1, respectively. *fimU* promoter activity of a *pilY1* mutant expressing each MP *in trans*. Expression of FimU, PilV, PilW, PilX, or PilE in the *pilY1* background had no impact on *fimU* promoter activity relative to the *pilY1* + empty vector control. (C) *fimU* promoter activity of *pilW* and *pilX* mutants overexpressing PilY1. Overexpression of PilY1 had no impact on *fimU* promoter activity in *pilW* and *pilX* backgrounds relative to the respective vector-only controls. Assays in (A), (B), and (C) were carried out in the presence of 0.05% L-arabinose to induce expression of the pBADGr promoter, n = 3.

**Fig S5.**
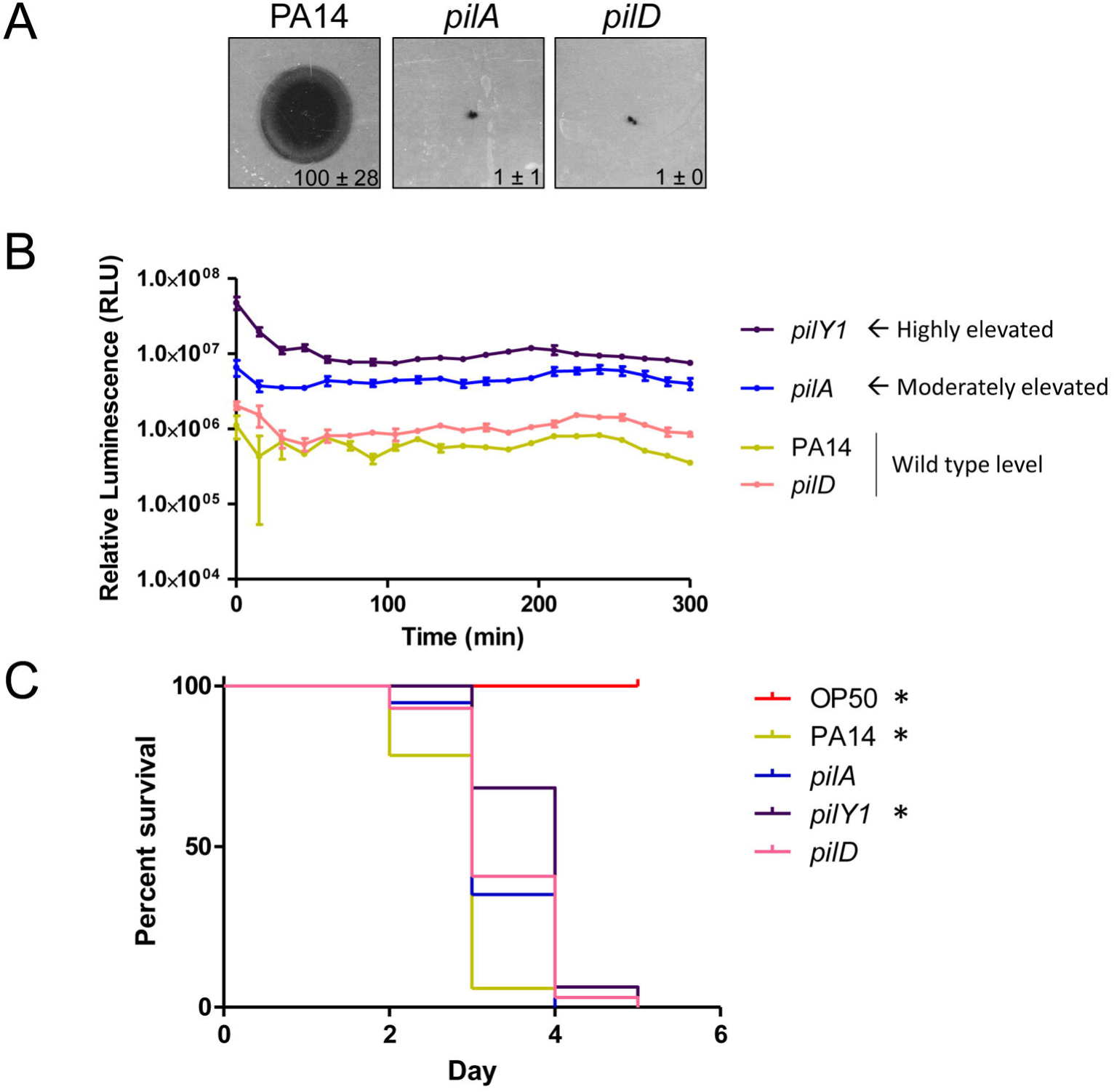
PilD is not required for PilWXY1-mediated modulation of FimS-AlgR activity. (A) Twitching motility assays for PA14 *pilA* and *pilD* mutants. Loss of *pilD* resulted in loss of twitching motility. Numbers indicate percent twitching area relative to WT, n = 3. (B) *fimU* promoter activity of a *pilD* mutant compared to PA14, *pilA*, and *pilY1*. Loss of *pilD* had no impact on *fimU* promoter activity relative to WT, n = 3. (C) SK assays for PA14, *pilA*, *pilY1*, and *pilD* mutants. A *pilD* mutant had equivalent virulence to a *pilA* mutant; less pathogenic than WT but more pathogenic than a *pilY1* mutants. Asterisks represent strains that were significantly different from the *pilA* mutant by Gehan-Breslow-Wilcoxon test at p = 0.05 (p = 0.0125 with a Bonferroni correction), n = 3.

**Fig S6.**
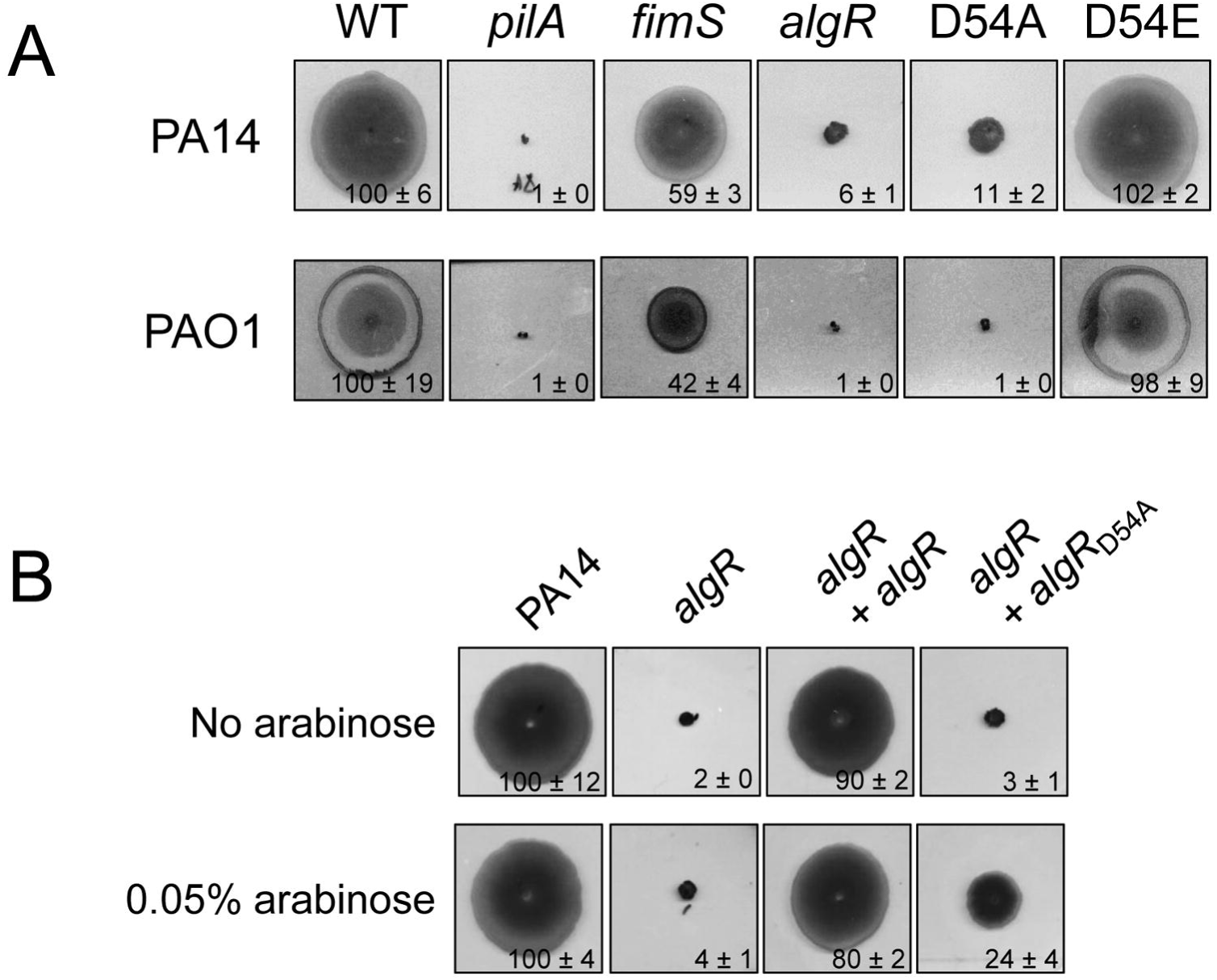
Phosphorylation of AlgR is required for optimal twitching motility. Twitching motility assays for PA14 *pilA*, *fimS*, *algR*, *algR*_D54A_, and *algR*_D54E_ mutants. Twitching motility was abolished in *pilA*, *algR*, and *algR*_D54A_ mutants, and fully retained in the *algR*_D54E_ mutant. A *fimS* mutant twitched to ~50% WT levels. (B) Twitching motility assays for PA14 *algR* complemented with AlgR or AlgR_D54A_. An *algR* mutant was fully complemented by AlgR with and without induction by 0.05% L-arabinose. The AlgR_D54A_ variant supported twitching motility in the *algR* mutant background in the presence of 0.05% L-arabinose, to ~25% WT levels. In (A) and (B), numbers indicate percent twitching area relative to WT, n = 3.

**Table S1.**
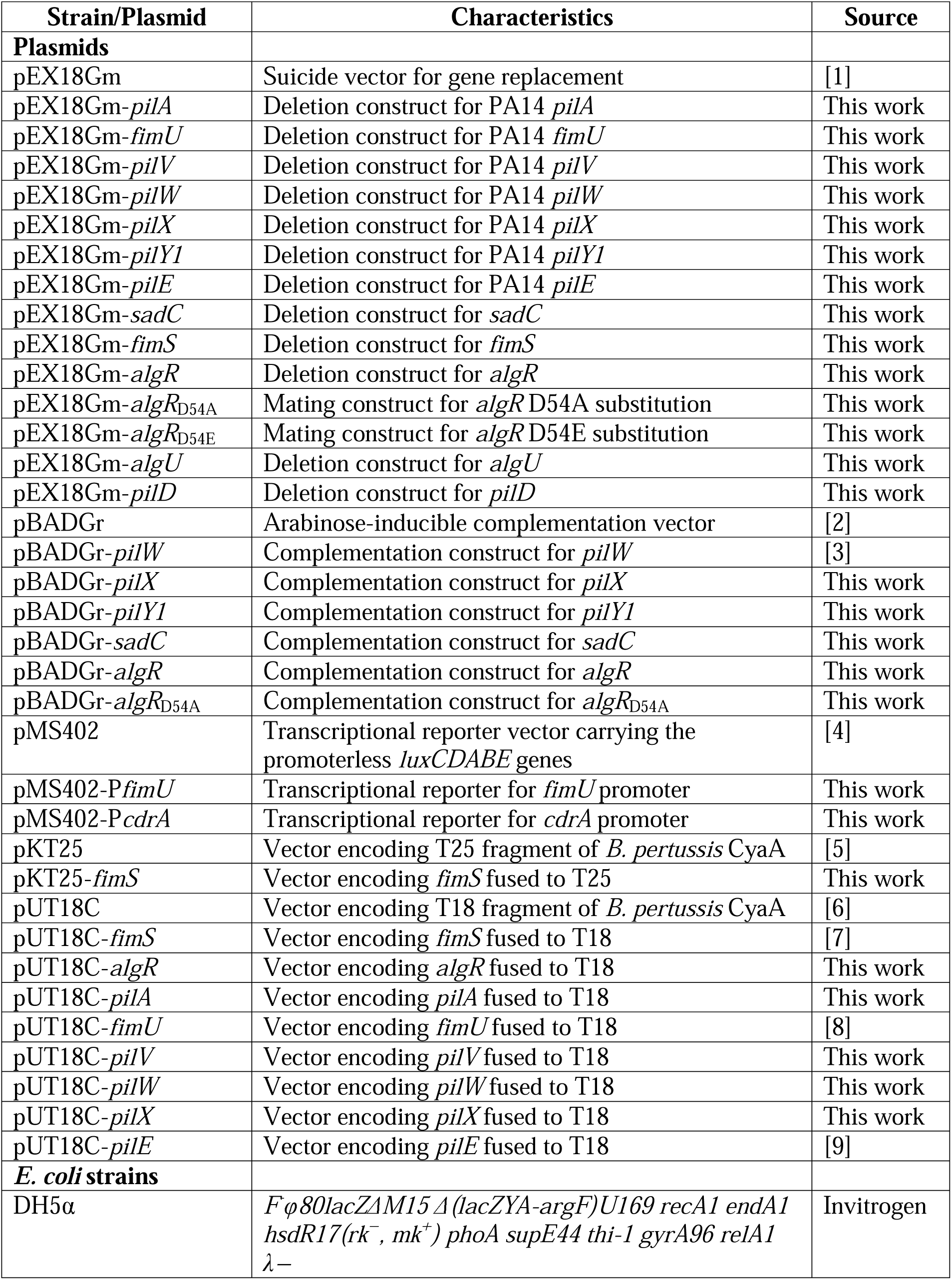
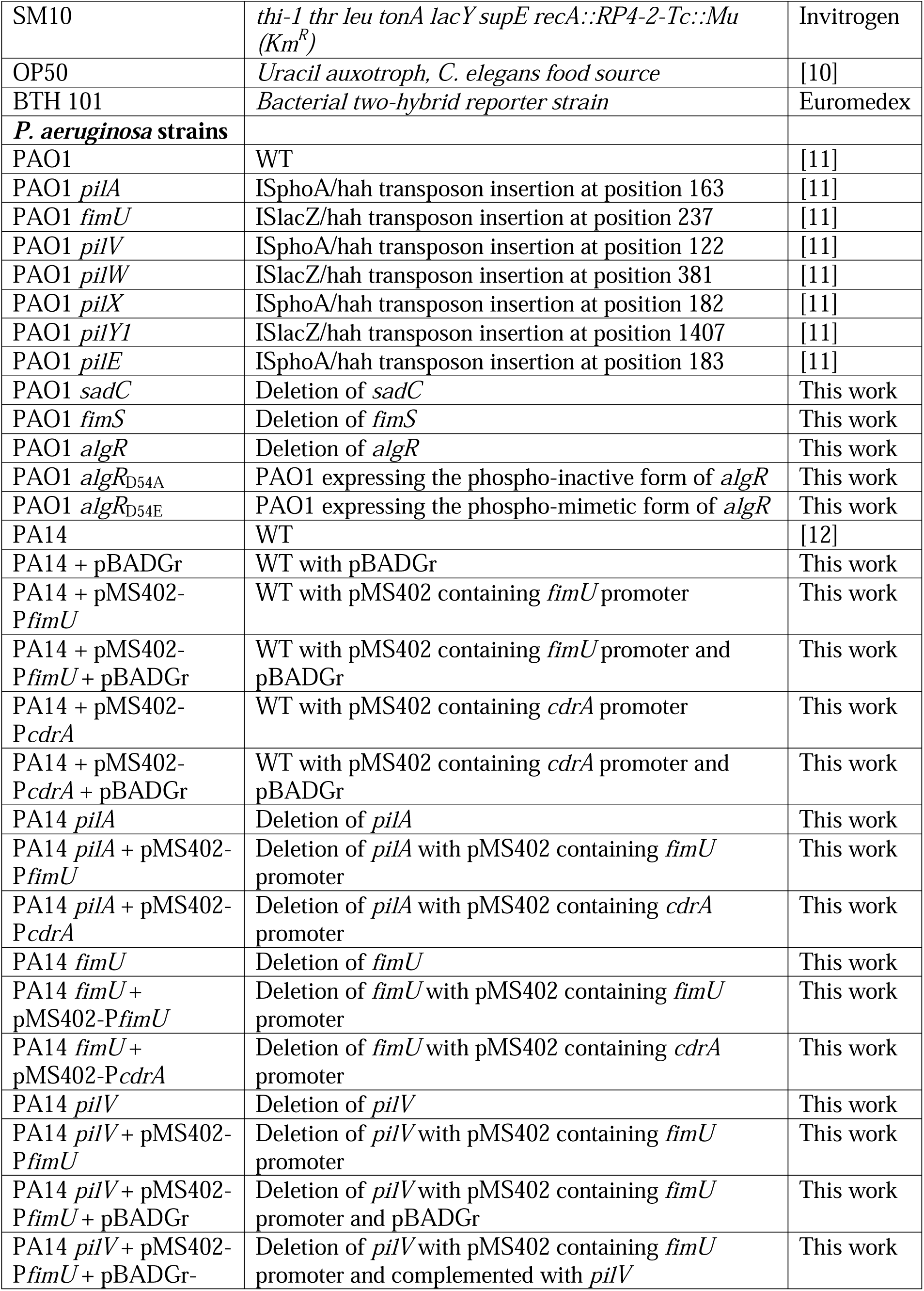
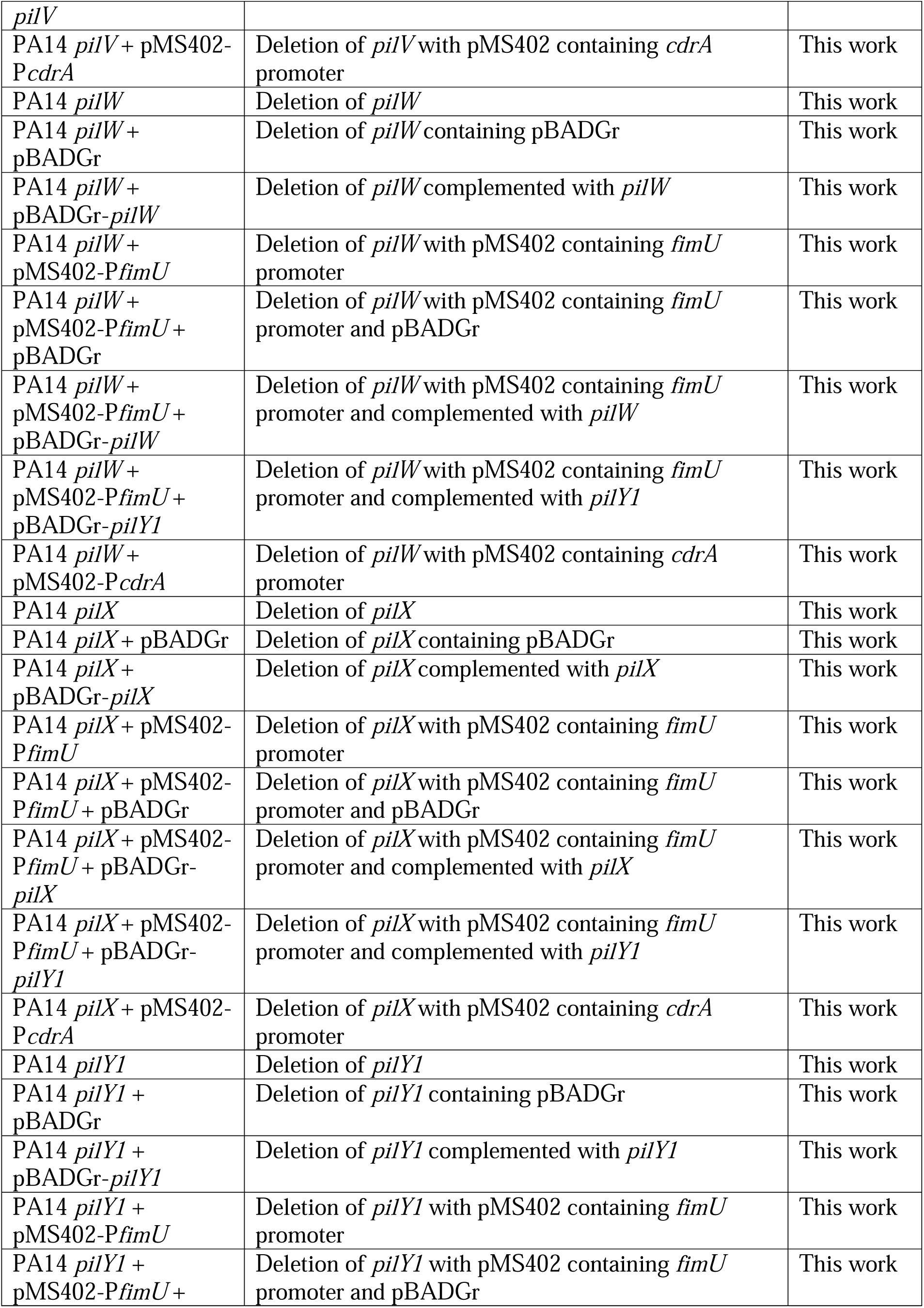
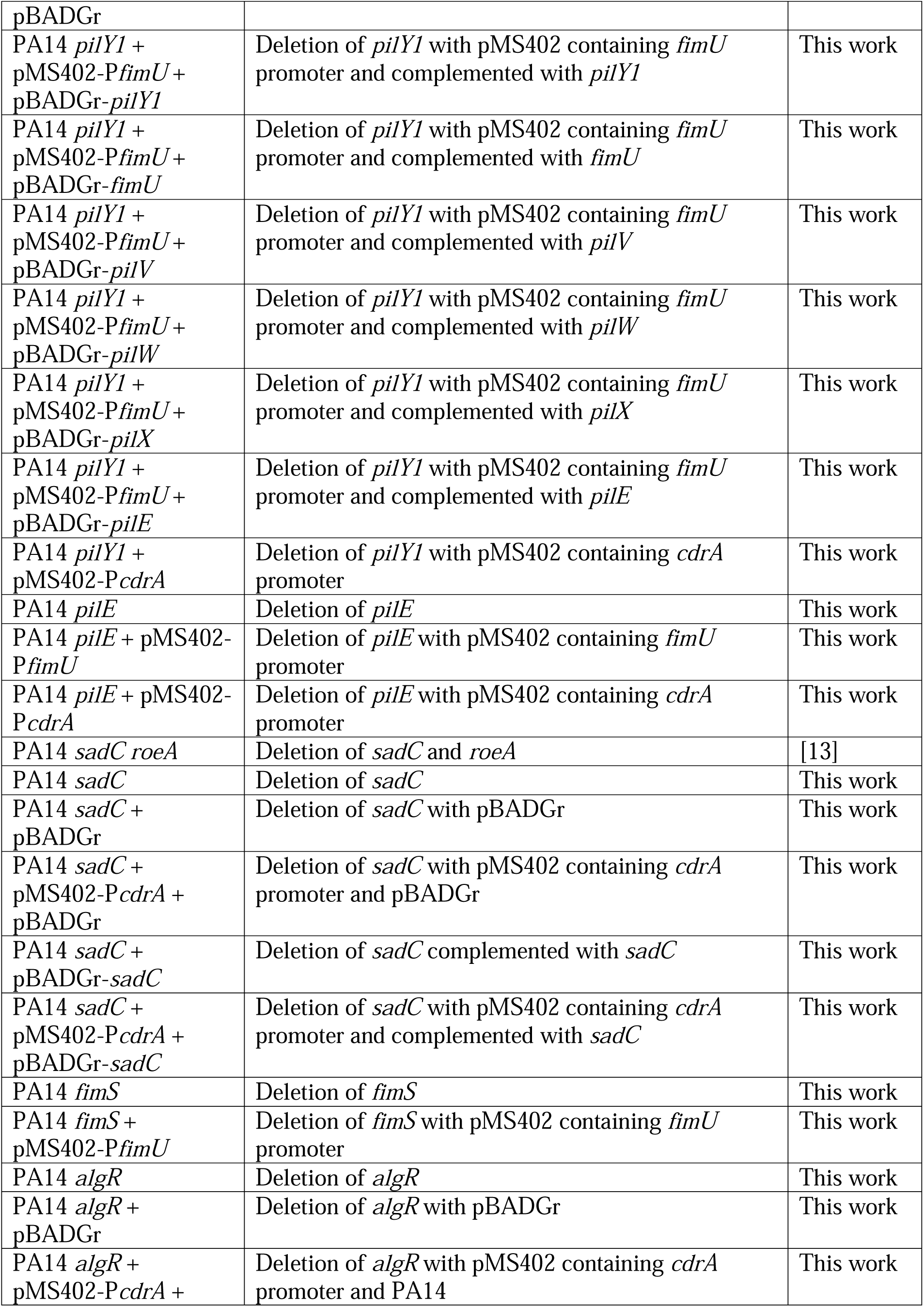
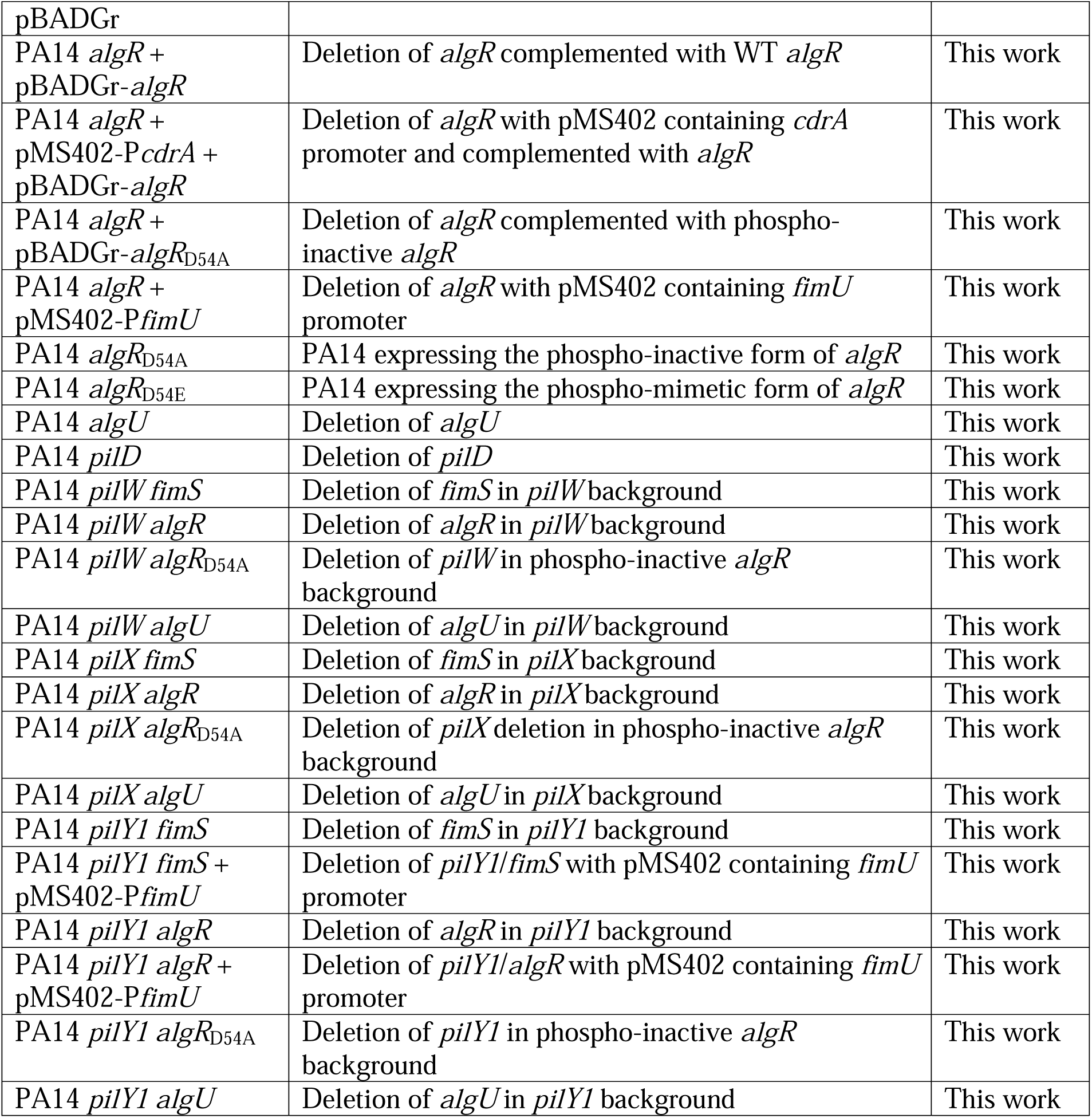
Bacterial strains and plasmids used in this study.

**Table S2.**
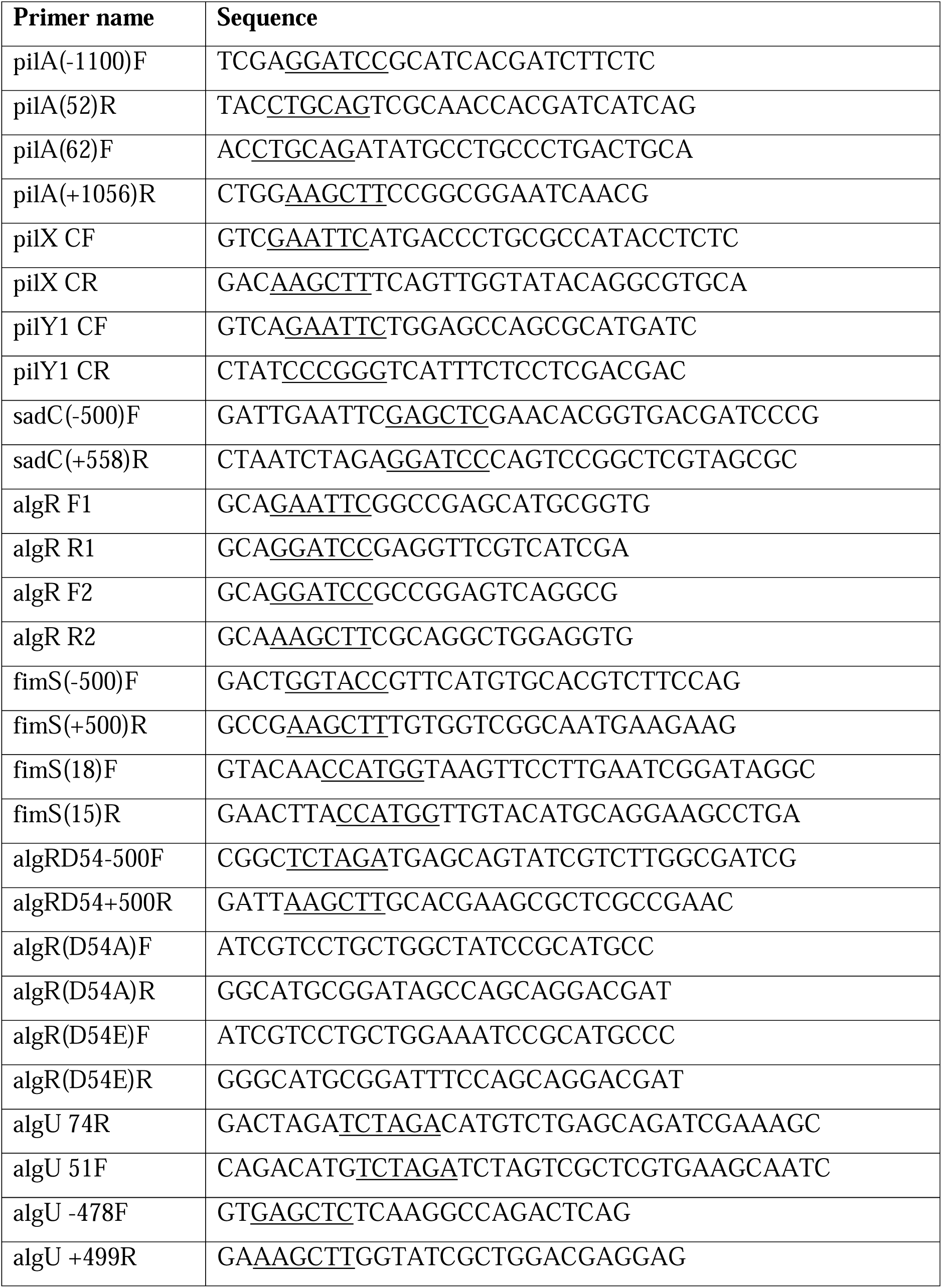
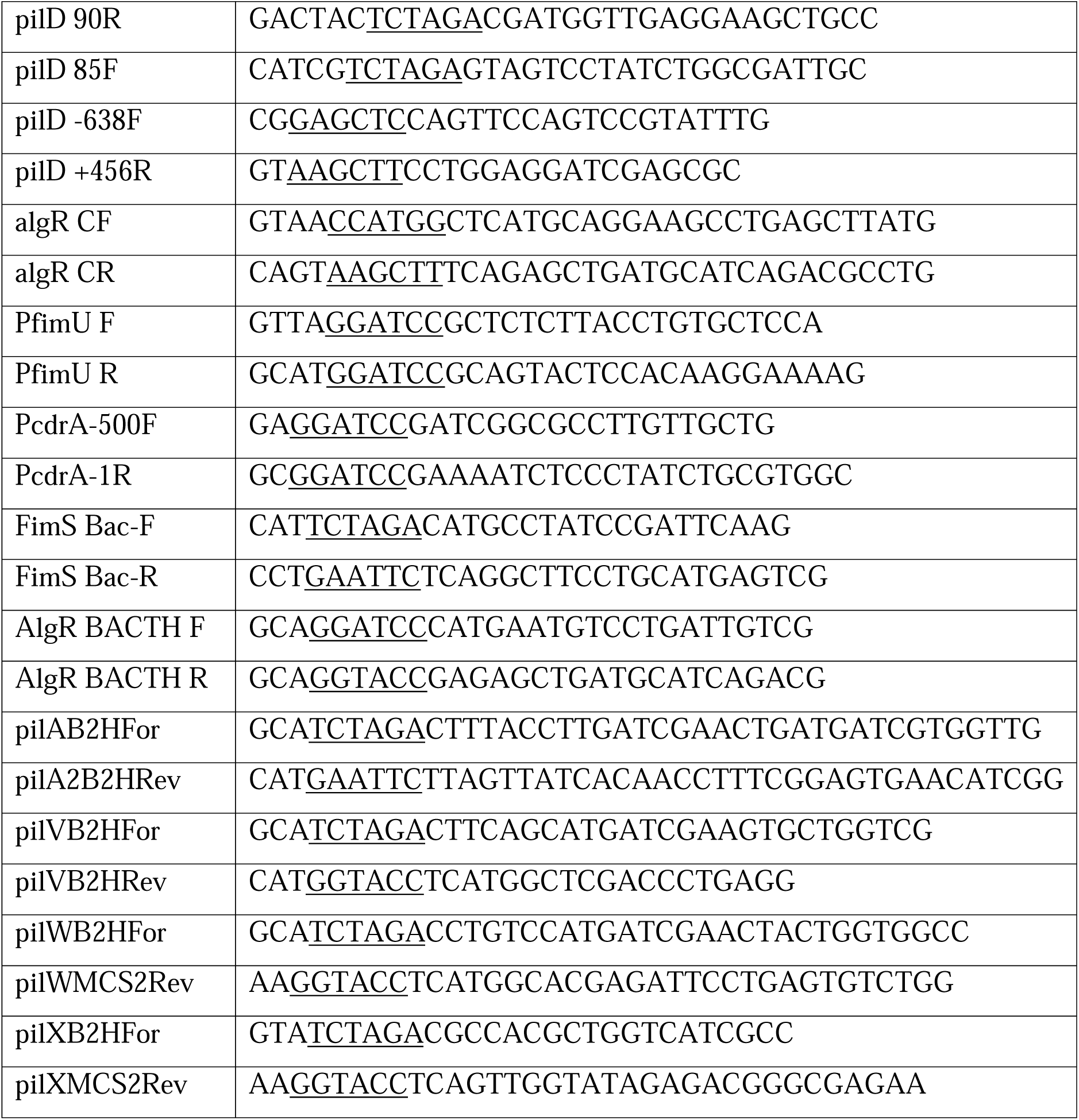
Primers used in this study. Restriction sites are underlined.

**File S1. Replicates for slow killing assays.** Three independent experiments for Figs 1A-B, 3B, 5A-B, 6A, 7A-C, 8, and Supplementary Figs S1B, S3A-B, S5C.

